# Enzymatic and Biophysical Analysis of two Highly Related Cytochrome P450 Reductases from *Artemisia annua* Reveals Differences in Their Ligand Interactions and Domain Motions

**DOI:** 10.64898/2026.05.13.725038

**Authors:** Bethany Mostert, Rika Judd, Thomas Makris, De-Yu Xie

**Affiliations:** Department of Plant and Microbial Biology, North Carolina State University, Raleigh, NC, USA; Department of Molecular and Structural Biochemistry, North Carolina State University, Raleigh, NC, USA

**Keywords:** Artemisinin, CPR, CYP, differential scanning fluorimetry, melting temperature

## Abstract

Artemisinin is an effective antimalarial drug sourced from *Artemisia annua,* but its low and variable yields require enhancement either semi-synthetically or *in-planta* to meet the global demand for treatment. Though essential enzymes have been identified in the artemisinin biosynthetic pathway, including an essential Cytochrome P450 monooxygenase (CYP71AV1), there are still many unknowns. Cytochrome P450 reductase 1 (herein, AaCPR1), has been experimentally confirmed as an electron transfer partner for CYP71AV1 in its three step oxygenation of key artemisinin precursors. However, the recent discovery of a highly related CPR, herein AaCPR2, introduces the possibility that another, potentially more catalytically favourable interaction, could exist for CYP71AV1. Therefore, enzyme kinetics and differential scanning fluorimetry (DSF) were used in the characterisation of both AaCPR1 and AaCPR2 to determine the existence and source of their catalytic differences. Tested enzyme activity under cytochrome *c* and NADPH concentrations revealed that AaCPR1 had lower K_m_ and higher k_cat_/K_m_ values, while AaCPR2 had higher V_max_ and k_cat_ values. This suggests that AaCPR1 is more effective at reducing cytochrome *c* when substrate conditions are limiting, whereas AaCPR2 is more effective than AaCPR1 at reducing cytochrome *c* when substrate conditions are saturating. This implies a functional partitioning of the two enzymes on the basis of substrate availability. The DSF results provided deeper insight into the different protein-ligand interactions between the two enzymes. AaCPR2 reached lower maximum melting temperatures across all tested conditions, whereas AaCPR1 had higher maximum melting temperatures. Thus, AaCPR1 exhibits higher thermal stability and has a higher temperature threshold than AaCPR2. This contributes to the notion that the AaCPRs are functionally divergent also on the basis of temperature. The cumulative differences in melting behaviour between the two enzymes led to the hypothesis that AaCPR1 and AaCPR2 exhibit different domain motions that may lead to preferential catalysis for one redox partner over another. This was further supported by the prediction of a highly variable loop region between the two enzymes at the connecting domain just after the flexible hinge. If such loops are highly mobile, as predicted, then the residue differences therein could provide a bio-structural basis for the kinetic and thermal/biophysical differences observed between AaCPR1 and AaCPR2. These data support that AaCPR1 and AaCPR2 possess fundamental biophysical differences despite their high degree of relatedness. Ultimately, these differences suggest differential metabolic functions of the two enzyme in artemisinin biosynthesis and/or other important secondary metabolic processes.

## Introduction

*Artemisia annua* is an effective antimalarial plant that produces artemisinin, an essential drug in Artemisinin based Combination Therapies (ACT) for the treatment of malaria (WHO, 2025). It is especially effective against cases caused by *Plasmodium* species such as *P. falciparum* that remain resistant to other antimalarials (Xia et al., 2025). The most recent report from the World Health Organization documented 282 million malaria cases that resulted in 610,000 deaths in 2024 (WHO, 2025). Producing sufficient amounts of artemisinin is one of the main approaches for reducing falciparum related deaths. In addition to its antimalarial capacities, artemisinin has been reported to possess other medicinal properties. Reports from preclinical studies have shown its involvement in anti-tumor activity (e.g., for breast, lung and colorectal cancers) and shown therapeutic potential in immune related diseases such as lupus, psoriasis, and Type 1 diabetes (Xia et al., 2025).

Artemisinin is an endoperoxide sesquiterpene lactone that is mainly isolated from the leaves of *A. annua*. To meet the global demand for ACT, approximately 25,000 tonnes of cultivated *A. annua* leaves are required to produce sufficient artemisinin (PATH, 2025). This large volume is necessitated by the varying content of artemisinin across plant populations (approximately 0.08-1.5% of dry weight for cross-pollinated commercial varieties) (Ikram & Simonsen, 2017; Weathers et al., 2006; Wetzstein et al., 2018). Attempts to overcome these low yields range from improved cultivation strategies and advanced breeding techniques, to developments in biotechnology, such as metabolic engineering (Kalalagh et al., 2025; Guo et al., 2023; Paponov et al., 2025; Wetzstein et al., 2018), and synthetic biology in microbial hosts (Paddon et al., 2013; Ro et al., 2008; Zeinali et al., 2025). These strategies have shown promising results with potential yield improvements, but continuous efforts are needed to make a significant impact.

These past efforts have made intensive progress in elucidating the artemisinin biosynthetic pathway. Presently, the steps from farnesyl diphosphate to artemisinic acid have been molecularly and biochemically characterized in the glandular trichomes of *A. annua* (Fig. 1). This involves the following enzymes: amorpha-4, 11-diene synthase (ADS), cytochrome P450 monooxygenase 71AV1 (CYP71AV1) (Teoh et al., 2006) and its partner cytochrome P450 reductase (CPR1) (Ro et al., 2008), artemisinic aldehyde reductase (DBR2) (Zhang et al., 2008), aldehyde dehydrogenase (ALDH1) (Teoh et al., 2009), and alcohol dehydrogenase (ADH1) (Paddon et al., 2013) (Fig. 1).

**Figure 1.**
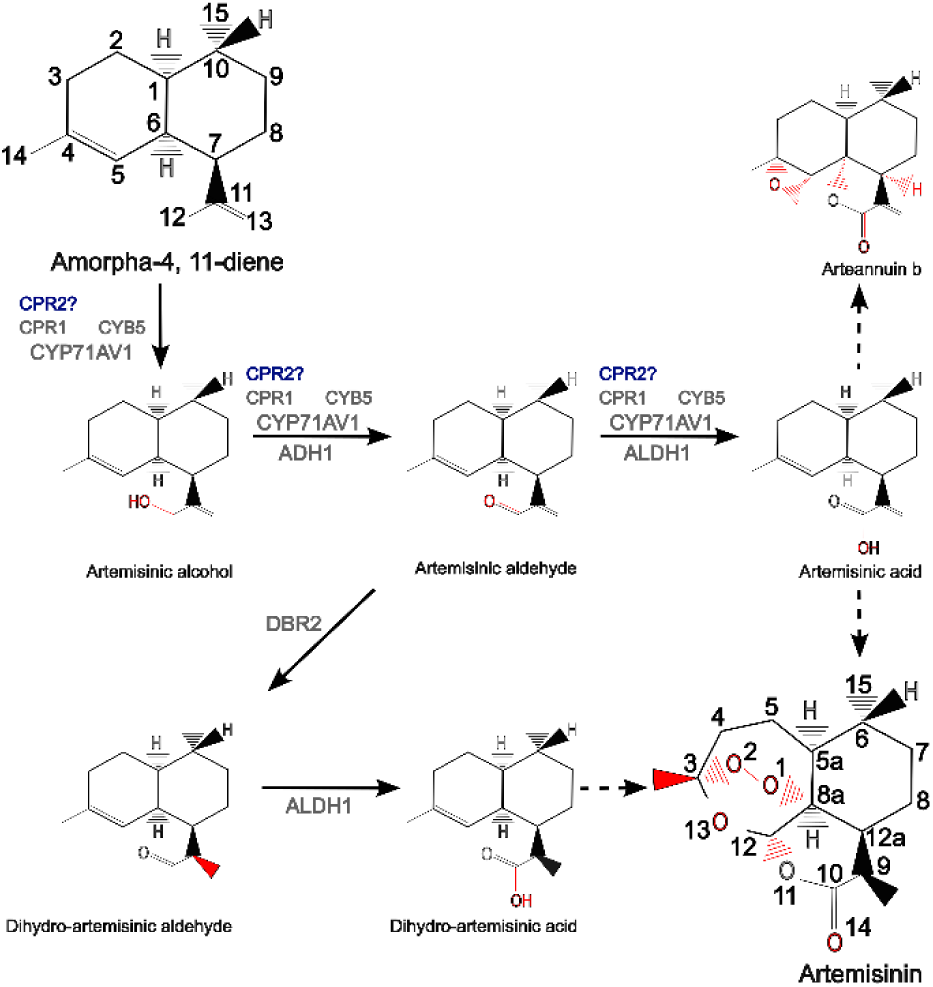
Figure 1. Biosynthetic pathway of Artemisinin from Amorpha-4, 11-diene. CYP71AV1 and its associated reductase (CPR1) are implicated in three major catalytic steps from amorpha-4, 11-diene to artemisinic acid. Solid arrows represent confirmed enzymatic reactions and dashed arrows represent non-enzymatic/unknown enzyme reactions. Abbreviated enzymes: Cytochrome P450 71AV1 (CYP71AV1), Cytochrome P450 reductase 1 (CPR1), Cytochrome b_5_ (CYB5), Cytochrome P450 reductase 2 (CPR2), Artemisinic alcohol dehydrogenase (ADH1), Artemisinic aldehyde Δ11(13) reductase (DBR2), Artemisinic aldehyde dehydrogenase (ALDH1). CPR2 is hypothesized to be involved in the CYP71AV1 enzymatic reactions and question marks (?) mean that its functions remain unknown. Furthermore, while A. annua CYB5 expression in a recombinant yeast strain is known to increase artemisinic alcohol content (Paddon et al., 2013), whether it interacts with it has not CYP71AV1 in planta remains unclear.

Among these characterised enzymes, the CYP71AV1-CPR1 partnership has been found to catalyze sequential steps in the pathway. Apart from catalyzing amorpha 4, 11-diene (the first artemisinin precursor) into artemisinic alcohol, these enzymatic partners are capable of performing additional oxidations that generate the subsequent aldehyde and acid (Teoh et al., 2006) (Fig. 1). In addition to CPR1, other reductase candidates have been proposed to interact with CYP71AV1 (Ma et al., 2015; Paddon et al., 2013). Paddon et al. (2013), for instance, suggested the participation of cytochrome b5 (CYB5), because its co-expression with CYP71AV1 and CPR1 in engineered yeast strains enhanced the production of artemisinic aldehyde. This suggests that additional oxidoreduction partners might be necessary to perform catalysis *in planta*. More recently, a second cytochrome P450 reductase 2 (CPR2) was genomically and transcriptionally verified in *A. annua* (Ma et al., 2015). Transcriptional analysis showed that the expression of CPR2 was closely associated with the presence of artemisinin in the leaves and flowers (Ma et al., 2015). Altogether, these support the hypothesis that CYP71AV1 may form complexes with more reductases than previously reported.

Cytochrome P450 reductases (CPRs) transfer electrons from NADPH through flavin cofactors to Cytochrome P450s (CYPs), cytochrome b5, and experimentally to nonphysiological redox partners such as cytochrome *c* and ferricyanide (Fukuchi-Mizutani et al., 1999; Huang et al., 2015). CPR is a small family. In humans and animals, it exists as only one gene. By contrast, two to four CPRs have been reported in plants and some fungi (Lah et al., 2008). Many plants possess at least two CPRs, which are divided into two phylogenetic clades known as CPR1 and CPR2. CPR1 is characterised to be widely associated with primary metabolism, while CPR2 is associated with secondary metabolism and differential expression under plant stress (Istiandari et al., 2023; K. Jensen & Møller, 2010; Urban et al., 1997). Though only a few CPRs from plants have been characterised (Choi et al., 2023; Istiandari et al., 2023; Jennewein et al., 2005; Laursen et al., 2016; Liao et al., 2024; Parage et al., 2016; Park et al., 2013; Qu et al., 2015; Ro et al., 2002; Rosco et al., 1997; Su et al., 2017; Yang et al., 2010; Zhang et al., 2022), previous research has shown that CPR isoforms exhibit diverging catalytic activities and enzyme associations. For instance, a co-expression of three CPR2 isoforms from *Oryza sativa* (rice) in *E. coli* showed that one of them consistently demonstrated higher catalytic activity with two cytochrome P450 enzymes: cinnamate 4-hydroxylase (*p-*coumaric acid biosynthesis) and tryptamine 5-hydroxylase (serotonin biosynthesis) (Park et al., 2013). A similar observation was made in the dicot species *Tripterygium wilfordii* (thunder god vine), which demonstrated that one out of four closely related CPRs exhibited preferential catalysis towards *ent*-kaurene oxidase (CYP701). In addition to preferential CYP catalysis, CPR isoforms are implicated in multi-enzyme complexes. Notably, a multi-protein complex consisting of three CPR2 isoforms—CPR2a-c, and key dhurrin biosynthesis enzymes CYP79A1, CYP71E1, were isolated from a lipid bilayer extraction of etiolated *Sorghum bicolor* (sorghum) seedlings (Laursen et al., 2016). This data suggests that even as CPRs diverge in their CYP preferences, they may also act in cohesion to synergistically support CYP catalyzed metabolism.

Like the mammalian CPR, plant CPRs possess essential functional domains for the transfer of electrons from NADPH to the heme of CYP. These domains include an N-terminal anchor for binding to the endoplasmic reticulum, a flavin adenine dinucleotide (FAD) binding domain, a flavin mononucleotide (FMN) binding domain, and an NADPH binding domain (Djordjevic et al., 1995; Niu et al., 2017). Numerous crystallisation and structural studies have been done on yeast, human, and rat CPR, alongside their various mutants, to elucidate mechanisms behind the interactions of these domains and their effects on the biophysical behaviour of CPRs (Burris-Hiday & Scott, 2024; Hamdane et al., 2009; Hubbard et al., 2001; Lamb et al., 2006; Wang et al., 1997; Zhao et al., 1996). In comparison, only ATR2 from Arabidopsis (Niu et al., 2017) and a CPR2 isoform (SbCPR2b) from *Sorghum bicolor* (Zhang et al., 2022) have been crystallised to understand these domains in plants. In agreement with rat and yeast CPRs, the crystals of ATR2 and SbCPR2b were captured in their closed states—a conformation that enables internal FMN-FAD electron transfer (Laursen et al., 2011; Niu et al., 2017; Zhang et al., 2022). Currently no crystals of plant CPRs in their open conformation have been deduced. Therefore, most plant CPR models rely on homology modeling with an open rat CPR ([3ES9.pdb](https://www.rcsb.org/structure/3ES9)), a feat only achieved by deleting four amino acids at the hinge region between the FAD and FMN binding groups (Hamdane et al., 2009). Mutation experiments and biochemical analyses have shown that this hinge region is at the crux of CPR conformational changes due to its inherent flexibility (Laursen et al., 2011; Campelo et al., 2018). Mutations and sequence differences within and near the CPR hinge have been widely investigated among rat and human CPRs for identifying key residues associated with open, closed, and intermediate conformational states of CPRs, which have effects on protein thermodynamics, ionic strength, and kinetics (Campelo et al., 2017, 2018; Esteves et al., 2020; Grunau et al., 2007). Functionally, this affects CPR interactions in a CYP and substrate dependent manner. Apart from the hinge region, other residues localised between binding domains affect the open and closed states of CPR. For instance, ATR2 contains an arginine residue (R708) at the salt bridge between the FMN and NADPH binding domains that is shared among plant and algae CPRs but not observed among mammals or yeast (Niu et al., 2017). When this residue was mutated, reductase activity to cytochrome *c* was abolished, leading to the hypothesis that R708 is an essential residue ‘switch’ that triggers the open and closed states of ATR2 in a plant specific manner (Niu et al. 2017).

Despite the plethora of research into the functional changes associated with residue mutations in human CPR, there is very little work done on how amino acid differences at these important inter-domain regions affect functions of plant CPR isoforms. Therefore, investigation on their mechanistic differences and behaviours at a biophysical and structural level are essential to comprehensively characterise the function of plant CPRs in plant metabolism and development.

As mentioned above, *AaCPR2* has been reported to express exclusively in *A. annua* leaves and flowers where artemisinin biosynthesis is most prevalent (Ma et al., 2015). Although the role of AaCPR2 in this antimalarial plant is unknown, the available transcriptional data support the hypothesis that AaCPR2 is involved in the biosynthetic pathway of artemisinin (Fig. 1). To test this hypothesis, recombinant AaCPR2 was purified in this work for biochemical and biophysical characterisation alongside AaCPR1 for comparative analysis. A protocol was developed to purify recombinant truncated AaCPR1 and AaCPR2 proteins induced in *E. coli*. Cytochrome *c* and both NADPH and NADH were used as substrates and co-enzymes for catalytic assays. Differential scanning fluorimetry (DSF) was performed to characterise the thermostability of the two CPR proteins in their free and ligand bound states. The results revealed different unfolding properties between the two enzymes, which suggests that residue differences between the two isoforms are associated with their unique structural and conformational behaviours. These data appear to be the first reported use of DSF to characterise the thermal unfolding properties of two plant CPR isoforms. Furthermore, these findings provide a solid foundation for immediate next metabolic characterisation of AaCPR2 in the biosynthesis of artemisinin.

## Results

### Phylogeny of AaCPR1 and AaCPR2 and Comparison of Their Conserved Domains

The amino acid sequences of both AaCPR1 ([ABM88789.1](https://www.ncbi.nlm.nih.gov/protein/124020687)) and AaCPR2 (A0A2U1KZS6.1) were accessed from the NCBI protein database. A comparison of their amino acid sequences yielded a 70.8% amino acid identity, with the major difference occurring at the N-terminal anchor, a region that has been reported to possess low conservation among other homologs (Cheng et al., 2023; Parage et al., 2016; Ro et al., 2002). The reductase features of AaCPR2 were confirmed through alignments to the empirically validated AaCPR1 (Simtchouk et al., 2013; Teoh et al., 2006) and ATR2 from *Arabidopsis thaliana*. The resulting alignment revealed that, like AaCPR1 and ATR2, the sequence of AaCPR2 contains the conserved FAD, FMN, and NADPH binding motifs (Fig. S1). Identity of the FMN-FAD hinge was evaluated from the tertiary protein structure as predicted by ColabFold (Mirdita et al., 2022) and found to be conserved among the ATR2 and two AaCPRs. The highest degree of variability between AaCPR1 and AaCPR2 is found within the connecting domain that proceeds from the hinge, bridging the FMN and FAD/NADPH domains.

These three sequences and 30 additional plant CPR sequences were aligned and used to generate a phylogenetic tree (Fig. 2). This tree grouped the CPR homologs into four clades: CPR1, CPR2, non-seed plant, and non-flowering plant groups. Both CPR1 and CPR2 clades were composed entirely of angiosperms. Most importantly, the tree grouped AaCPR1 and AaCPR2 into the CPR2 clade (Fig. 1). This observation is reflective of the high sequence similarity of these enzymes. Additionally, two CPRs listed as CPR1 and CPR2 from angiosperm species *Aconitum vilmorinianum* and *Tripterygium wilfordii* grouped within the CPR2 clade (Cheng et al., 2023; Su et al., 2017). This, along with the presence of multiple CPR2 isoforms among the grasses (e.g., *Zea mays, Oryza sativa*), suggests that AaCPR1 and AaCPR2 may in fact be two CPR2 isoforms in *A. annua*. Based on these findings, it would be more accurate to refer to AaCPR1 as AaCPR2a, and AaCPR2 as AaCPR2b, as reported among other CPR2 isoforms (Park, Kim, Sanjeewa, et al., 2013; Zhang et al., 2022). This proposed new annotation is further supported by the longer amino acid sequences (when compared to CPR1) and serine and threonine enriched N-termini of the AaCPRs (visible in the alignment, Fig. S1), which is consistent with reports comparing CPR1 and CPR2 enzymes (Parage et al., 2016; Ro et al., 2002).

**Figure 2.**
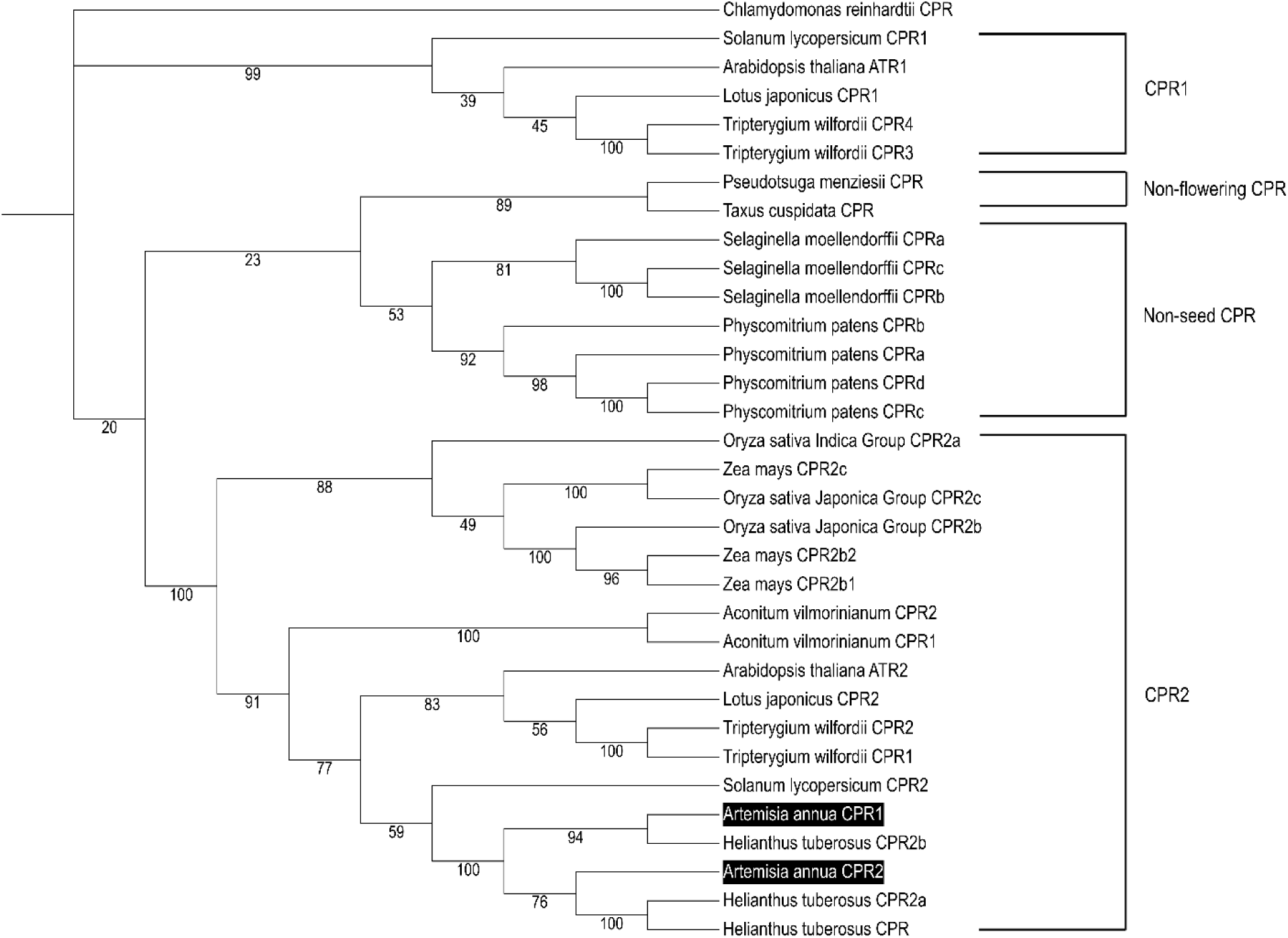
An unrooted phylogenetic tree of CPR developed from thirty-three CPR amino acid sequences. This phylogenetic tree was constructed using the maximum likelihood method with [Mega11](https://www.megasoftware.net/) (Tamura et al., 2021), which was further annotated with [ITOL](https://itol.embl.de/). *Chlamydomonas reinhardtii* (algae) CPR was used as an outgroup. Node values represent the percentage of supporting bootstrap replicates out of 1000 iterations. The resulting tree groups CPRs into the four following clades: CPR1, non-flowering CPRs, non-seed CPRs, and CPR2. AaCPR1 and AaCPR2 are grouped in the same (CPR2) clade.

### Establishment of Protocols for the Purification of Soluble Truncated AaCPR1 and AaCPR2

The full-length gene sequences for *AaCPR1* (<u>EF197890.1</u>) and *AaCPR2* (contig sequence from Ma et al., 2015) were cloned using the primers listed in Table S1. Based on predictions with the [TMHMMserver](https://services.healthtech.dtu.dk/services/DeepTMHMM-1.0/) (v. 2.0), transmembrane domains of the two AaCPRs at their N-termini were identified and then removed through PCR to obtain truncated *Δ1-66 AaCPR1* and *Δ1-67 AaCPR2* fragments. After purification of these PCR products, *Δ1-66 AaCPR1* and *Δ1-67 AaCPR2* were cloned into His-tagged Gateway vectors pDEST17 (Fig. 3A), and HisMBP-pDEST17 (Fig. 4A). The resulting recombinant plasmids were introduced into *E. coli* for protein induction (Fig. 3, Fig. 4). Given that the complete oxidation of the flavin groups produces a yellow colour (Simtchouk et al., 2013; Zhang et al., 2022), highly expressing suspension cultures resulted in orange cell pellets (Fig 4E). The resulting purified protein samples for AaCPR1 and AaCPR2 also exhibited a yellow colour (Fig. 3D and Fig. 4E).

**Figure 3.**
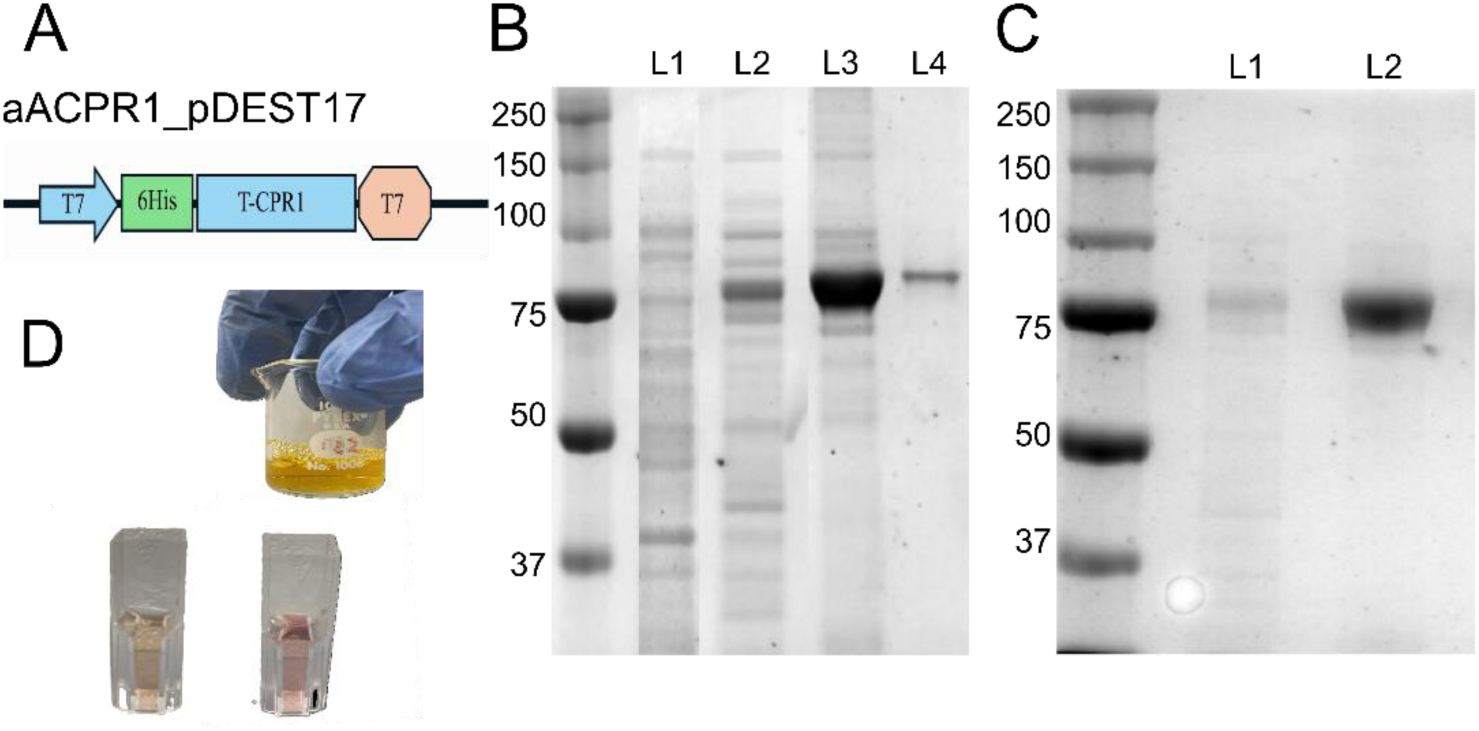
**A scheme showing the expression induction and purification of recombinant AaCPR1 from E. coli**. A. Gateway expression vector pDEST17 was used to induce AaCPR1 after truncation of the first 66 amino acids and fused to a His-tag at the N-terminus. B. An image of SDS/PAGE shows a purified AaCPR1 via a Nickel (Ni-NTA) column. Molecular weight markers are in kDa. L1, crude protein extract from empty vector as control; L2, crude protein extract; L3-L4, eluted AaCPR1. C. An image of SDS/PAGE shows purified AaCPR1 via AKTA chromatography. AaCPR1 elution from the nickel purification were combined and run on a HiTrap IMAC high performance column. L1, crude cell lysate; L2, AKTA purified AaCPR1. D. Images show the bright yellow colour of purified AaCPR1 and the reduction of Cytochrome c from orange to pink) by AaCPR1.

**Figure 4.**
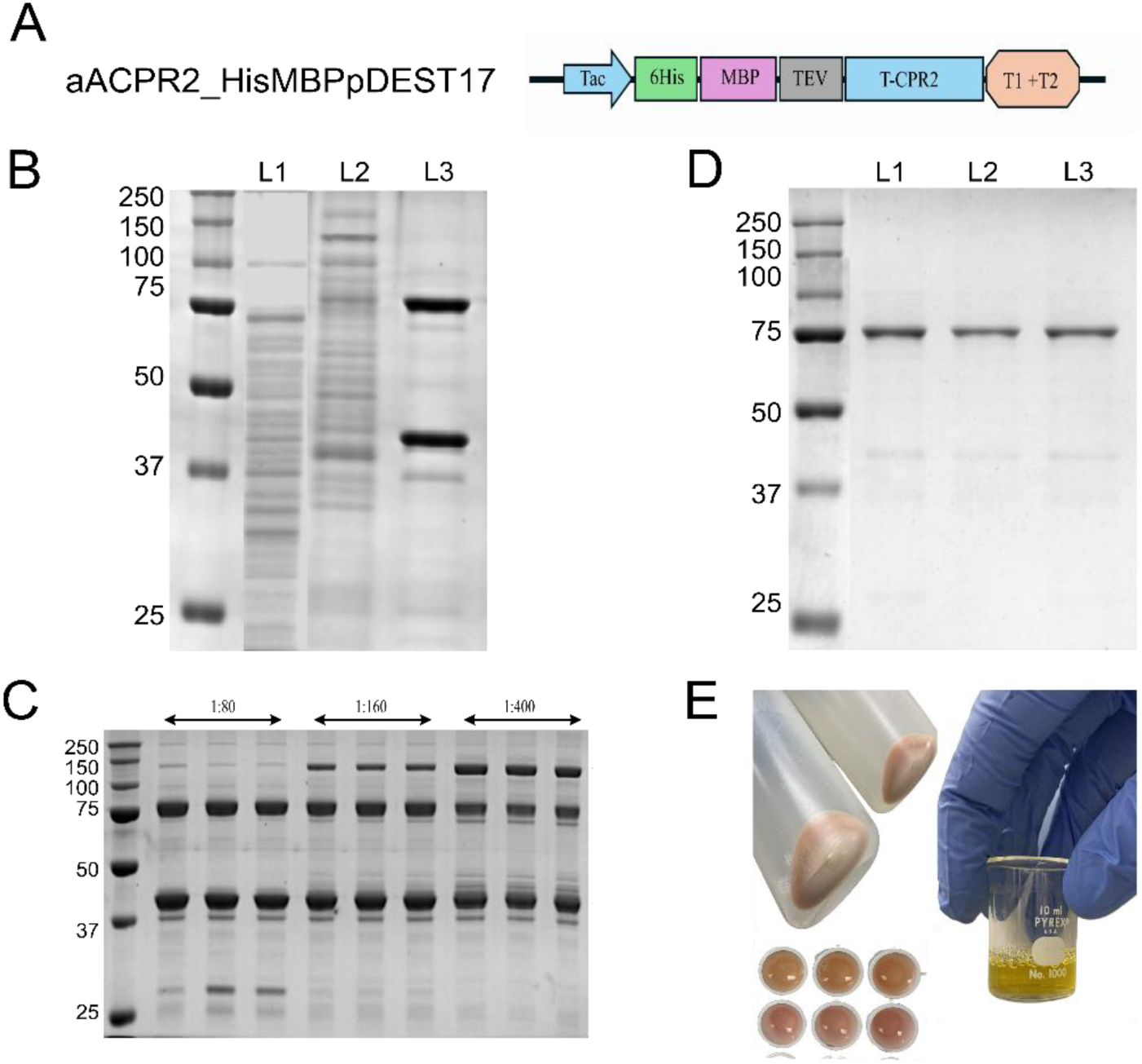
A scheme showing expression induction and purification of recombinant AaCPR2 from E. coli A. **A.** Gateway expression vector His-MBP pDEST17 was used to express AaCPR2 after truncation of the first 67 amino acids and fused to a His-MBP tag at the N-terminus **B. An image of** SDS/PAGE shows AaCPR2 purified via Nickel (Ni-NTA) and amylose columns. Molecular weight markers are in kDa. L1, crude protein extract from empty vector as control; L2, crude protein extract. The 118 kDa protein band between 150 and 100 kDa is the MBP-AaCPR2 fusion protein); L3, two strong bands resulted from the TEV digestion of the MBP-AaCPR2 fusion protein purified via an Ni-NTA column, the 75 kDa being AaCPR2 and the 42kD band being MBP. **C. An image of** SDS/PAGE shows that the 1:80 ratio of TEV to MBP-AaCPR2 effectively digested fused protein to 75 kDa AaCPR2 and 42 kDa MBP, while 1: 160 and 1: 400 ratios performed partial digestion. Each ratio was completed with three replicates. **D. An image of** SDS/PAGE shows 75 kDa AaCPR2 purified from TEV digestion of MBP-AaCPR2 via an amylose (MBP-binding) column. L1-L3, three purification elutions. **E.** Images show yellowish colour of AaCPR2 induced in E. coli cells (bright orange) and purified from amylose columns (bright yellow colour) and pinkish colour from the reduction of Cytochrome c by AaCPR2.

This phenotype greatly enhanced purification efforts as protein purification was visually monitored in both the Ni-NTA and MBP affinity amylose columns that were used. SDS/PAGE analysis showed that both *Δ1-66 AaCPR1* and *Δ1-67 AaCPR2* expressed recombinant AaCPR1 and AaCPR2. The molecular weight (MW) of AaCPR1 was approximately 75kDa (His-tagged amino-acid sequence: 71.8kDa) (Fig. 3B-C) and MBP-AaCPR2 was approximately 120kDa (His-MBP tagged sequence with TEV cut site: 115.82kDa) (Fig. 4B-C).

Though MBP fused CPRs have been shown to establish Michaelis-Menten kinetics with cytochrome *c* (Su et al., 2017), the MBP tag in this study was removed prior to downstream biochemical characterizations to prevent potential non-native protein-protein interactions or MBP fusion associated issues. After the initial pass through the Ni-NTA column, the purified MBP-tagged AaCPR2 was further digested with Ice-TEV protease to cleave the His-MBP tag. SDS-PAGE analysis showed that the digestion produced tag-free, truncated AaCPR2 with an approximate 75kDa band (truncated amino-acid sequence is 71.87kDa) (Fig.4D). To further assess the quality of the TEV digestion and MBP removal method, a Western blot using a polyclonal MBP antibody was performed on the MBP-AaCPR2 fusion protein and the MBP-free AaCPR2 (Fig. S2). The MBP antibody did not bind to the MBP-free AaCPR2, confirming the absence of MBP on the truncated protein due to successful digestion. Both the truncated AaCPR1 and AaCPR2 were incubated with cytochrome *c*, a natural electron acceptor to assess their reduction capabilities. The incubations showed a colour transition from an orange to pink (Fig. 3D and Fig. 4E), validating their electron transfer capacities. These results demonstrate that the induction and purification protocols were effective in isolating soluble and active truncated AaCPR1 and AaCPR2 for enzymatic characterization.

### Spectral Characteristics of AaCPR1 and AaCPR2

The cytochrome P450 reductase identity of both truncated AaCPR1 and AaCPR2 was further confirmed through spectral analysis in the 200-750 nm region. The reduction activity of the enzymes was observed through anaerobic incubation of 50µM enzyme with 0-100µM NADPH, to create different molar ratios (R_NADPH_) of NADPH:CPR (Fig. 5). The recorded UV-visible absorbance spectral profiles revealed that the spectral profiles of both AaCPR1 and AaCPR2 in their fully oxidized forms without NADPH were composed of two notable peaks at 380 nm and 454 nm, a shoulder at 474 nm, and little to no absorption beyond 470 nm. These features are consistent with those observed among other diflavins (Simtchouk et al., 2013; Zhang et al., 2022). The effects of NADPH on the spectral profiles were analysed in detail. Upon its addition, the flavin absorbance of both the proteins between 300-470 nm began to drop and a peak emerged in the 500-650 nm range, which is characteristic of blue semiquinone spectra and partial reduction of the flavoprotein from hydride ion transfer (Aigrain et al., 2011; Murataliev et al., 2004) (Fig. 5). This absorbance trait is also consistent with the formation of NADP^+^, FADH^•^, and FMN^•-^, which are the result of interflavin electron transfer from FAD to FMN (Sevrioukova et al., 1996). The effect of NADPH on this absorption peak varied across its added concentrations. After the first addition of 25 μM NADPH (1:1 R_NADPH_), the semiquinone band stabilised and remained relatively unchanged. Thus, NADPH was unable to completely reduce the enzymes to their four electron states (FADH_2_ and FMNH_2_). For complete reduction/depletion of the FAD/FMN moieties, most spectral experiments use a molar excess of dithionite (Na_2_S_2_O_4_) (Simtchouk et al., 2013; Vigil et al., 2021). The peak that formed and gradually increased at 340 nm represents the absorbance spectrum of NADPH (De Ruyck, 2007) (Fig. 5).

**Figure 5.**
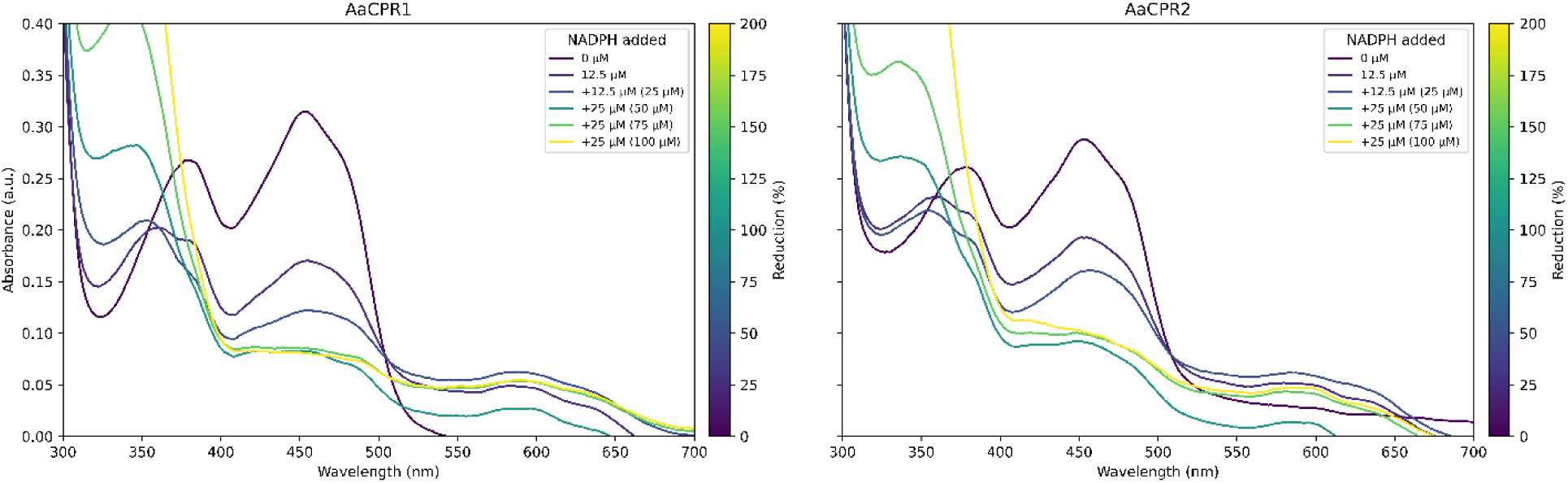
Reduction of flavin absorbance spectra of AaCPR1 and AaCPR2 by NADPH. The concentrations of AaCPR1 and AaCPR2 were 50μM in 50mM Tris-HCl (pH 7.5). The absorbance spectra of AaCPR1 and AaCPR2 were altered by NADPH under anaerobic conditions. Five concentrations of NADPH, 12.5μM, 12.5μM, 25μM, 25μM, and 25μM, were sequentially added to reach final 25μM, 50μM, 75μM and 100μM concentrations for the reduction of the two AaCPRs. The resulting five mixtures of NADPH were scanned from 325 nm to 700 nm. The spectral profiles were recorded to characterise the effects of NADPH concentrations on flavin absorption. The absorbance peaks correspond to characteristic flavin maxima.

### Kinetic Characterization of AaCPR1 and AaCPR2 with NADPH, NADH, and Cytochrome *c*

The kinetics of AaCPR1 and AaCPR2 were characterized with the substrate cytochrome *c* and two co-enzymes, NADPH and NADH. All experiments were completed using experimentally determined optimal conditions with 50mM Tris-HCl pH 7.5 buffer, at 25°C and 10nm of enzyme (see Table S2 for a complete list of tested buffers). To facilitate high throughput data for enzyme kinetics, 96 well plates were favoured over cuvettes and read in a UV-Vis plate reader. Fresh enzyme concentrations were replenished every three hours throughout the kinetic experiments, since thawed CPR lost catalytic activity over time. Enzyme samples were freshly prepared from the −80°C stocks and used to continue the experiment as needed. This ensured that the saturation of the CPRs was reflective of their true enzymatic behaviour rather than as an artifact of post-thaw enzyme instability. The changes in absorbance for all kinetic assays were measured at 550 nm based on the reduction of cytochrome *c.* The enzyme velocities were calculated using its extinction co-efficient (21/mM/cm) and then plotted (Guengerich et al., 2009). The resulting kinetic parameters (V_max_, K_m_, k_cat_, and K_m_/k_cat_) for cytochrome *c*, NADPH, and NADH are included in Table 1.

**Table 1.** Kinetic parameters of AaCPR1 and AaCPR2 for cytochrome *c* (0.25μM-50μM), NADPH (1μM-75/100μM) and NADH (100μM-50mM).

|  | Cytochrome c |  | NADPH |  | NADH |  |
| --- | --- | --- | --- | --- | --- | --- |
|  | AaCPR1 | AaCPR2 | AaCPR1 | AaCPR2 | AaCPR1 | AaCPR2 |
| $V_{\max}$ ($\mu\text{mol}/\text{min}^{-1}$) | $3.71 \pm 0.40$ | $5.08 \pm 0.59$ | $2.60 \pm 0.14$ | $4.00 \pm 0.39$ | $2.38 \pm 0.25$ | $1.01 \pm 0.16$ |
| $K_m$ ($\mu\text{mol}$) | $3.37 \pm 0.76$ | $5.9 \pm 1.4$ | $2.81 \pm 0.71$ | $7.2 \pm 1.8$ | $3848 \pm 1200$ | $3424 \pm 2000$ |
| $k_{\text{cat}}$ ($\text{min}^{-1}$) | $371 \pm 40$ | $508 \pm 69$ | $260 \pm 14$ | $400 \pm 39$ | $238 \pm 25$ | $101 \pm 16$ |
| $k_{\text{cat}}/K_m$ | $110 \pm 28$ | $87 \pm 24$ | $92 \pm 24$ | $56 \pm 15$ | $0.062 \pm 0.020$ | $0.029 \pm 0.018$ |

To characterise the kinetic parameters for cytochrome *c*, the enzymatic reactions were carried out across a substrate concentration from 0-75µM in the presence of 100μM NADPH. The resulting plots showed that the two enzymes followed typical Michael-Menten kinetics (Fig. 6 A-B).

**Figure 6.**
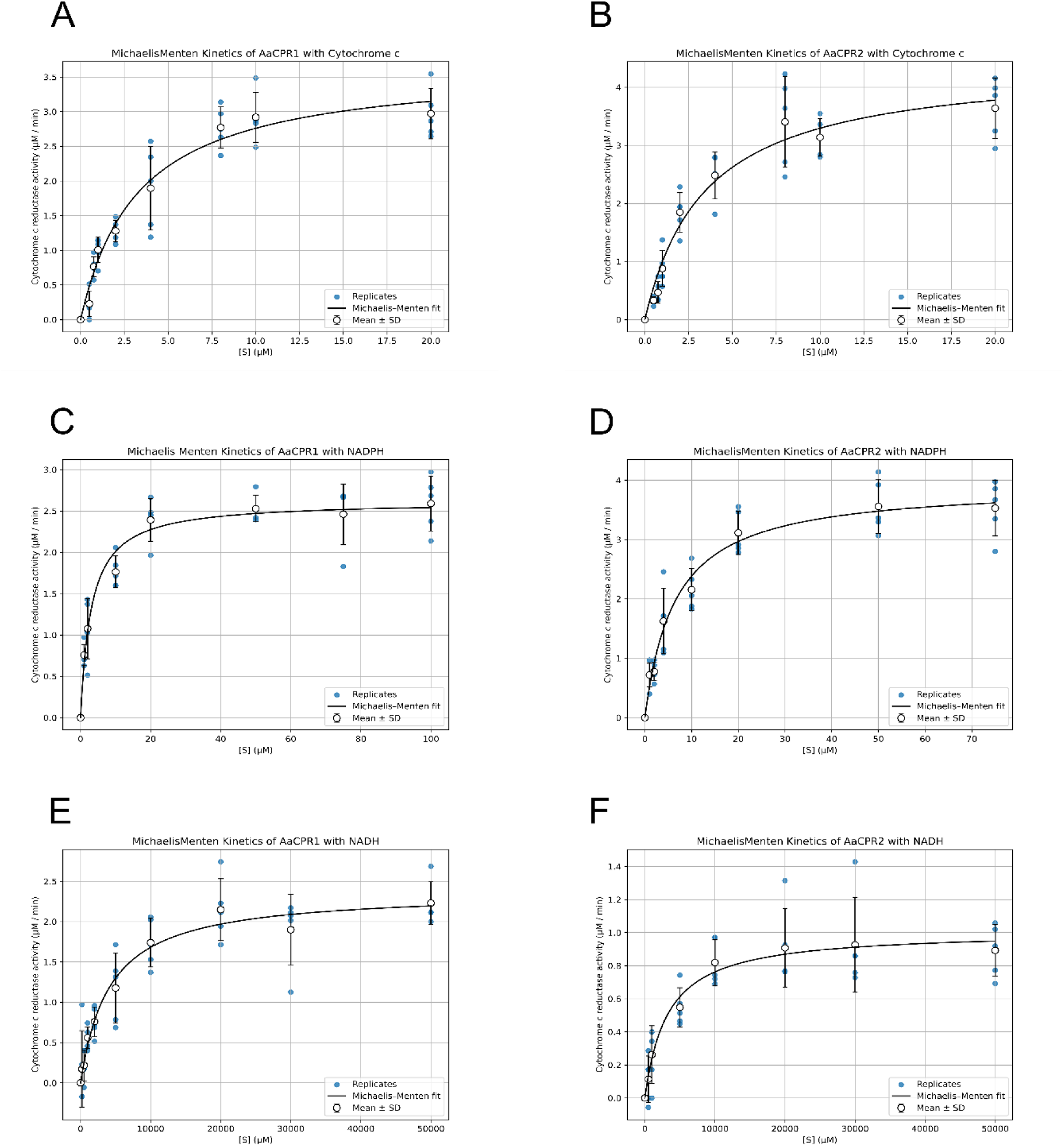
Plots showing Kinetic Curves for AaCPR1 and AaCPR2 against Cytochrome c, NADPH, and NADH. 10nm of AaCPR1 and AaCPR2 were used for each reaction. Catalysis followed Michaelis Menten kinetics with cytochrome *c* (0.25μM-50μM), NADPH (1μM-75/100μM) and NADH (100μM-50mM). Five replicates, represented by the blue dots, were included for calculating initial velocities at each substrate concentration. The line represents the average fit across the plotted velocities, denoted with open black circles and error bars (standard deviation). **A-B:** AaCPR1 (A) and AaCPR2 (B) against Cytochrome c. **C-D:** AaCPR1 (C) and AaCPR2 (D) against NADPH. **E-F:** AaCPR1 (E) and AaCPR2 (F) against NADH.

While other studies have reported that CPR purification can prevent FMN from properly loading onto the protein, both AaCPRs positively interacted with cytochrome *c,* validating that they had sufficient amounts of FMN to execute electron transfer (Fig. 6A, 6B) (Aigrain et al., 2011; Simtchouk et al., 2013). Moreover, the presence of sufficient FMN in both CPRs was also evident based on their reasonable k_cat_ values (Whitelaw et al., 2015). This is based on findings that k_cat_ values from FMN limited CPRs are sub-optimal (e.g., k_cat_ 0.12^-s^ from FMN depleted ATR1) (Whitelaw et al., 2015). The cytochrome *c* K_m_ values of both CPRs were similar to those of other plant CPRs (e.g., WsCPR1 (5.06 ± 0.30) and WsCPR2 (6.48 ± 0.33) from *Withania somnifera*) (Rana et al., 2013). Based on the K_m_ values, AaCPR2 has a lower affinity for cytochrome *c* than AaCPR1. However, the resulting V_max_ and K_cat_ values indicate that AaCPR2 has a higher turnover effectiveness to reduce cytochrome *c* than AaCPR1. To analyse kinetic parameters of NADPH, the enzymatic reactions were performed across a co-factor concentration from 0-100µM in the presence of 10μM of cytochrome *c*. The results showed that the two enzymes followed Michelis-Menten kinetics with NADPH (Fig. 6 C-D). The K_m_ values of NADPH for AaCPR1 and AaCPR2 were also similar to that of other reported plant CPRs, such as ATR2 (1.02 ± 0.012) (Whitelaw et al., 2015), GhCPR1 and GhCPR2 (4.5 ± 0.2; 5.6 ± 0.6) (Yang et al., 2010), and SbCPR2a (5.3 ± 0.5) and SbCPR2b (8.9 ± 1.0) from sorghum (Zhang et al., 2022). Once again, the K_m_ values revealed that AaCPR2 has a lower affinity for NADPH than AaCPR1; however, the V_max_ and K_cat_ values reveal a higher turnover efficiency to use NADPH for electron transfer than AaCPR1. When compared to NADPH, the high K_m_ values for AaCPR1 (3800 µM) and AaCPR2 (2500µM) against NADH (Table 1) indicate that it was a poor co-enzyme substitute. Therefore, like other plant CPRs, AaCPR1 and 2 appear to catalytically prefer NADPH (Simtchouk et al., 2013). Under all conditions, based on K_m_ and k_cat_ values, AaCPR1 displayed a higher overall enzyme efficiency.

The values were calculated using weighted non-linear regression against five replicates at each substrate concentration. Significant figures for kinetic parameters are dictated by the precision of the uncertainty (standard error, ±). A kinetic curve plotter tool, [EKViS](https://github.com/bhmoster/EKViS_Enzyme-Kinetic-Curve-Visualisation-Suite/tree/main), was developed for parameter extraction and plotting.

### Characterization of Melting/Unfolding Behaviour for AaCPR1 and AaCPR2 Under Different Ligand Conditions via Differential Scanning Fluorimetry Assay

Differential Scanning Fluorimetry (DSF) is a high throughput approach that is useful for detecting protein-ligand interactions (Bai et al., 2019). When an enzyme is stabilised by a ligand, this is reflected as an increase in the melting temperature of that enzyme—the protein unfolds and a fluorescent dye binds to its hydrophobic regions. An increased melting temperature indicates strong ligand binding, particularly as the concentration of the ligand increases (Bai et al., 2019). The DSF or Thermofluor assay was used to test the structural stability of both AaCPR1 and 2 under the following ligand conditions: a six condition ligand panel, increasing concentrations of NADPH with an appropriate concentration of cytochrome *c* (20µM), increasing concentrations of cytochrome *c* with an appropriate concentration of NADPH (30µM), increasing concentrations of NADPH alone, and increasing concentrations of cytochrome *c* alone. To ensure that the biphasic behaviour and unfolding responses of each experiment were a direct result of AaCPR activity and not as an artifact of cytochrome *c* unfolding responses, cytochrome *c* was included alone with the fluorescent dye. Cytochrome *c* alone demonstrated little to no fluorescence under the same condition; thus, it did not dictate the unique unfolding patterns observed among AaCPR1 and AaCPR2 in the DSF assay.

Under standard DSF analysis, such as CFX melt curve peak export data, replicate to replicate variation can cause meaningful biological peaks to be missed. To improve this analysis, the raw exported relative fluorescence units of AaCPR1 and 2 recorded for each experiment were used to develop a more sensitive peak calling tool in Python (DSF-PeakSpeak). The resulting derivative curves were used to characterise the dynamic unfolding of AaCPR1 and AaCPR2. In addition to each derivative plot, melting temperature tables were included to better capture multiple melting transitions, as the derivative curves in the plots resulted from average values and thus do not always visually represent the diverse unfolding behaviour seen among the replicate curves (see “ReplicateCurves_DSF.pdf” for a complete representation of replicate curves at all conditions). DSF-PeakSpeak was used to analyse and develop plots for all of the following DSF experiments.

A variety of ligands and ligand combinations were initially tested to differentiate the unfolding behaviour of the enzymes across these conditions (Fig. 7). AaCPR1 possessed a clear biphasic melting pattern as an apo protein (non-coenzyme bound), when bound to cytochrome *c,* and when combined with cytochrome *c* and NADPH (Fig. 7A). The average melting temperatures for the first and second transitions under these conditions were 47°C and 60°C, respectively. Interestingly, the peak depth of the second melting transition was similar for the no ligand control and NADPH alone, but it decreased under the cytochrome *c* and NADH conditions. This demonstrates more restrained unfolding behaviour for AaCPR1 when bound to NADH and cytochrome *c*. This interpretation is based on the quenched fluorescent signal under these conditions, which indicates that less dye was binding to hydrophobic regions of the protein. AaCPR2 exhibited different biphasic melting patterns (Fig. 7B), wherein the no-ligand, NADPH, and cytochrome *c* + NADPH conditions recorded two melting transitions at average temperatures of 44°C and 55°C (Fig. 7B).

**Figure 7.**
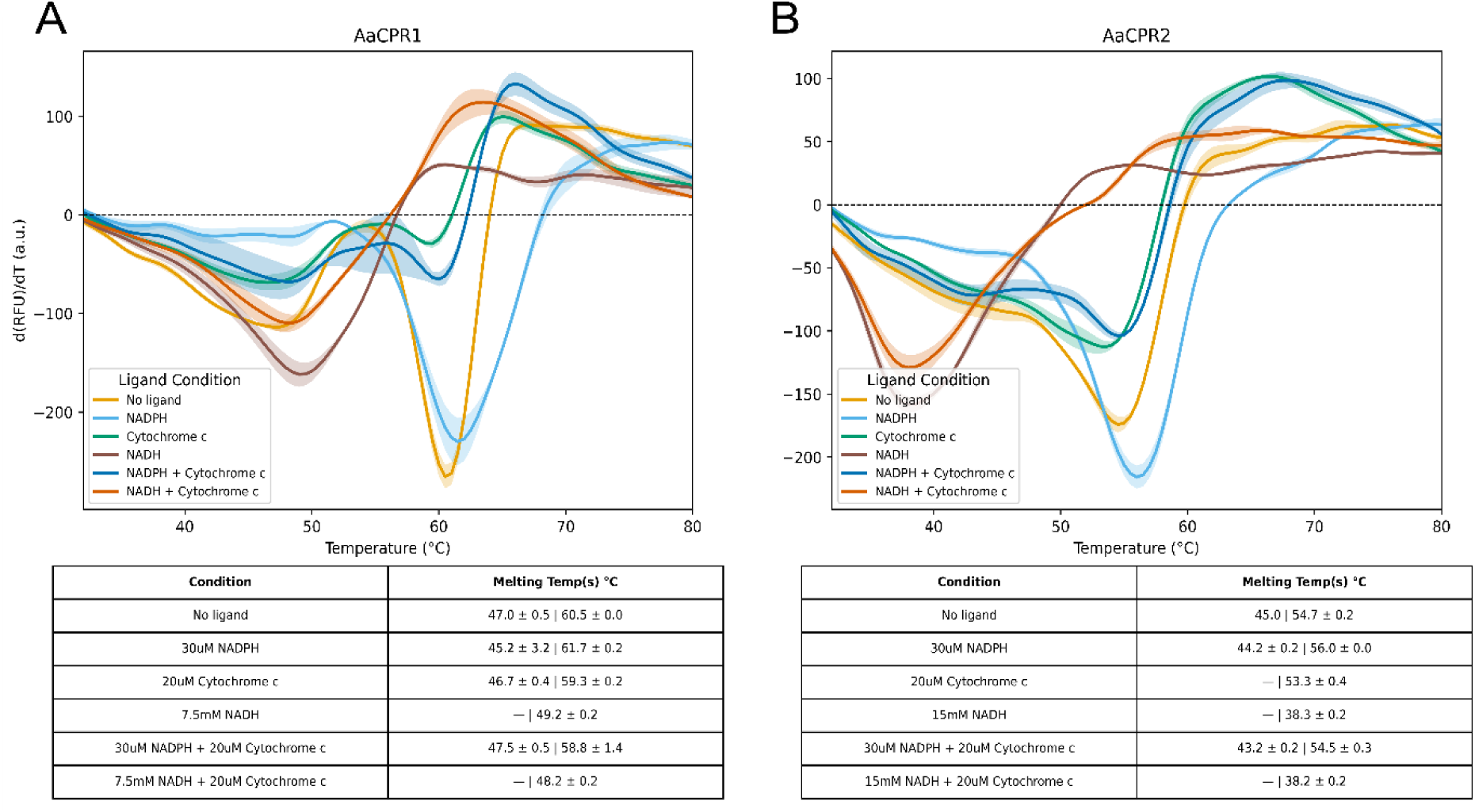
Effects of substrate ligands on AaCPR1 and AaCPR2 melting temperature (Tm) values. A differential scanning fluorimetry (DSF) assay was completed to measure Tm values for the two proteins with NADPH (30µM), NADH (7.5mM or 15mM), and cytochrome *c* (20µM) either alone or in combination with each other. Plots show the average derivative curves across 3 replicates for each condition. One lower and another higher melting temperature transition were detected for the two enzymes. **A.** NADH, cytochrome *c*, and their combination reduced the major Tm value of AaCPR1 while NADPH slightly increased it. **B.** NADH, cytochrome *c*, and their combination reduced the major Tm value of AaCPR2 while NADPH increased it.

The addition of NADPH (30µM) alone resulted in a positive melting shift for both the enzymes (Fig. 7 A-B). Cytochrome *c* (20µM) either alone or with NADPH slightly reduced the melting temperatures for AaCPR1, and AaCPR2. NADH (7.5mM and 15mM) either alone or with cytochrome *c*, eliminated a secondary melt curve entirely and dramatically decreased the melting temperature of both enzymes. Altogether, these data show different biphasic and monophasic melting features between AaCPR1 and AaCPR2 under various conditions.

### Effects of NADPH and Cytochrome *c* Concentrations on the Biphasic Behaviour of AaCPR1 and AaCPR2

Given the large drop in melting temperature when both enzymes were incubated with NADH and its poor kinetic performance, this ligand condition was left out of subsequent experiments. The effect of NADPH concentration on the two enzymes was next tested in association with 20µM cytochrome *c* (Fig. S3A). Interestingly, increasing NADPH (1-200μM) had no effect on the thermal shift of AaCPR1 when combined with cytochrome *c.* The second melting transition in particular lost energetic dominance (visualized as shallower peak depths) at higher concentrations of NADPH. While AaCPR2 exhibited a slight increase in the melting temperature for the second transition under higher NADPH concentrations, its effect was minimal, raising no more than 1.2°C above the no-ligand control. Of greater interest was that its peaks maintained their energetic dominance better than AaCPR1 despite increasing NADPH concentrations with cytochrome *c*.

The effect of cytochrome *c* on the two enzymes was then tested in the presence of 30μM NADPH (Fig. S3B). The tested concentrations of cytochrome *c* were 0.75-100μM. At lower concentrations of cytochrome *c* (0.75-2μM) with NADPH, both AaCPR1 and AaCPR2 slightly increased their melting temperature at their second transitions: AaCPR1 displayed increases from 0.7°C (2μM cytochrome *c*) to 1.7°C (0.75μM cytochrome *c*), and AaCPR2 displayed increases from 1.5°C (2μM cytochrome *c*) to as high as 6.2°C (0.75μM cytochrome *c*). Except for these concentrations, there appeared to be no major effect of increasing cytochrome *c* concentration on protein stability when NADPH was present. In fact, similar to the effect observed with NADPH concentrations and cytochrome *c* (Fig. S3A), when cytochrome *c* reached 20μM and above, the magnitude of the second transition decreased for AaCPR1 and AaCPR2 (Fig. S3A).

NADPH (1-200μM) (Fig. 8) and cytochrome *c* (0.75-100μM) (Fig. 9) alone were then tested to understand their effects on the two enzymes. Not only did increasing NADPH alone result in a monotonic melting temperature increase for AaCPR1, but it also altered its biphasic melting behaviour (Fig. 8A). As NADPH concentrations increased, the initial melting transition for AaCPR1 gradually lost energetic dominance, particularly at 30μM and above (Fig. 8A). In comparison, biphasic melting was more dynamic for AaCPR2 (Fig. 8B). The first and second melting transitions displayed similar energetic dominance, and in one experimental repeat, melded together to form one wide, unified melting peak (see ReplicateCurves_DSF.pdf). The first melting temperature was unstable across all concentrations of NADPH (1-200μM) (Fig. 8B). Comparatively, the second melting temperature of AaCPR2 stably increased from 53.8°C to 57.5°C across 1-50 μM NADPH, dropped to 57°C at 100μM NADPH and then increased to 58°C at 200μM NADPH (Fig. 8B). These data indicate that energetic dominance was more stochastic for CPR2 at no ligand, 100μM, and 200μM than at other concentrations of NADPH tested (Fig. 8B).

**Figure 8.**
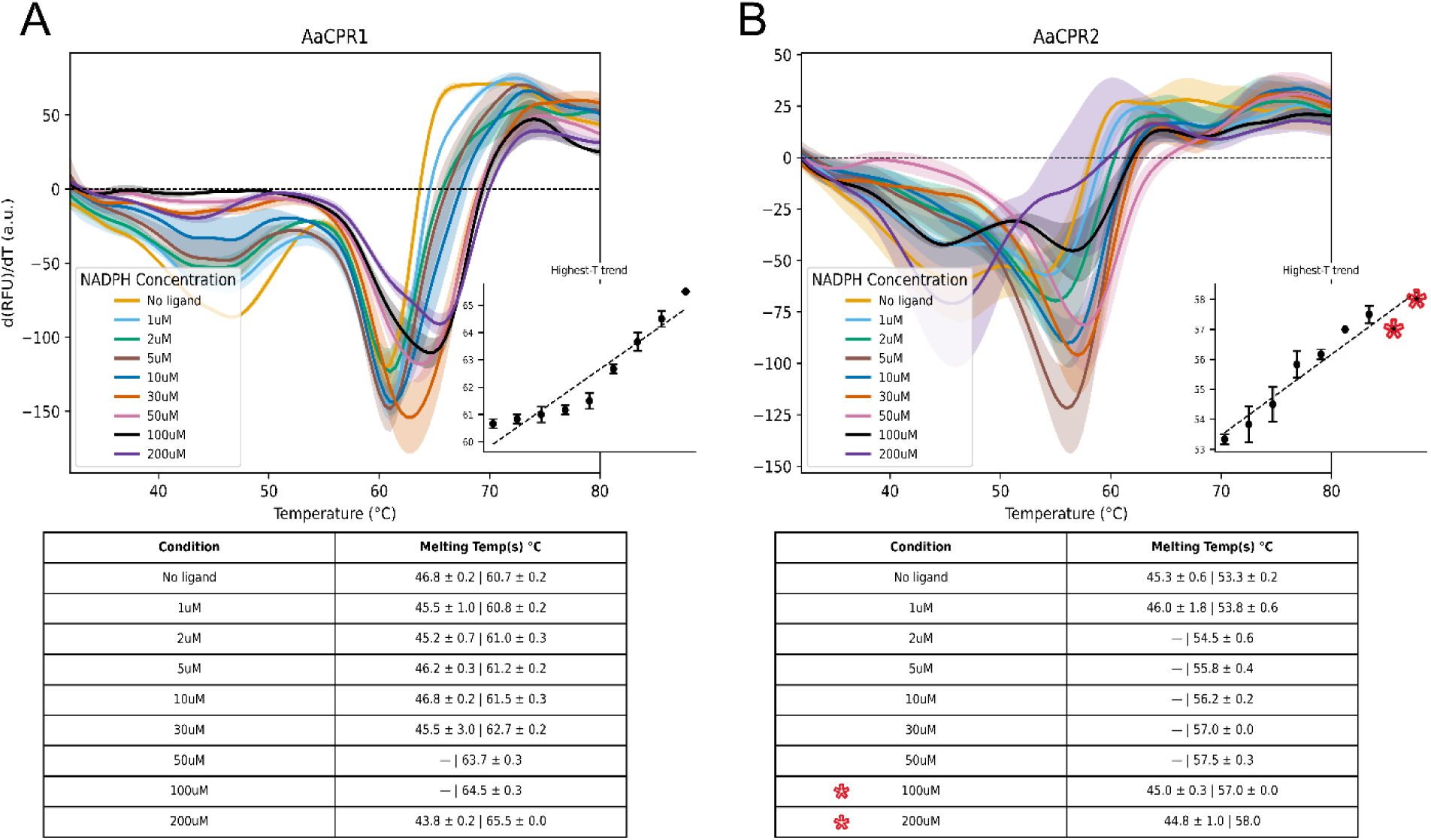
Effects of NADPH on AaCPR1 and AaCPR2 melting temperature (Tm) values. Eight concentrations of tested NADPH ranged from1uM-200uM in 50mM Tris-HCl (pH 7.5). Derivative curves were plotted for the two enzymes, and an insert showing the trend of the second Tm was included to visualize the linear regression of the highest recorded temperatures for AaCPR1 and AaCPR2 ± standard deviation. **A.** NADPH gradually increased the dominant second Tm value of AaCPR1 **B.** AaCPR2 lost the first melting transition from 2-50μM and was then regained at 100µM and 200µM (highlighted by red stars). A gradual increase was observed for the second Tm as NADPH concentrations increased. At 100 and 200uM of NADPH, the Tm of AaCPR2 became dynamic.

**Figure 9.**
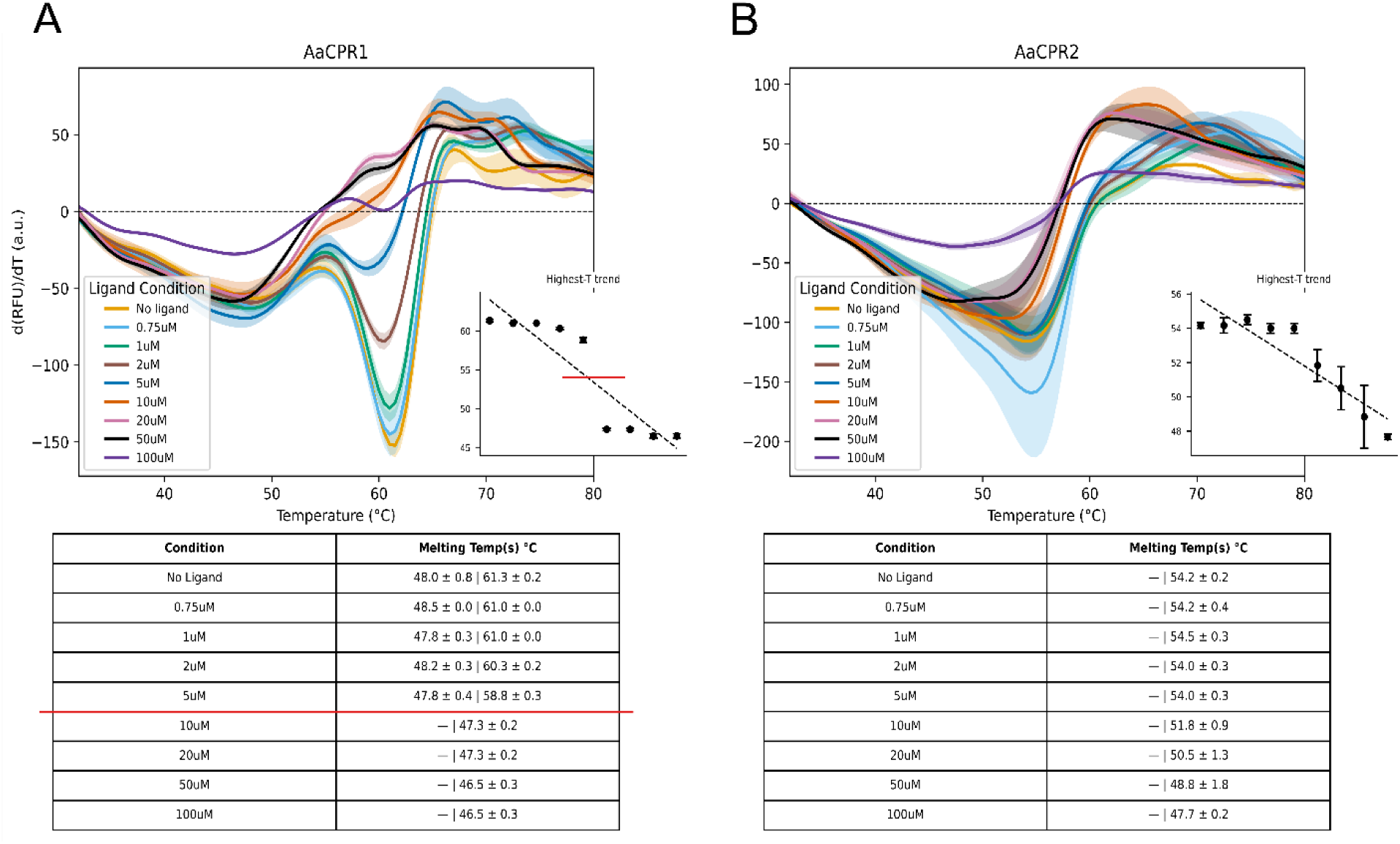
Effects of Cytochrome c on AaCPR1 and AaCPR2 melting temperature (Tm) values. Eight concentrations of tested cytochrome *c* ranged from 0.75-100μM in 50mM Tris-HCl (pH 7.5). Derivative curves were plotted for the two enzymes, and a temperature trend insert was included to visualize the linear regression of the highest recorded temperatures for AaCPR1 and AaCPR2 ± standard deviation. **A.** The first melting transition for AaCPR1 gradually lost energetic dominance as cytochrome *c* concentrations increased. The second melting transition gradually decreased from 0.75 to 5µM and was then abruptly lost after 10µM of cytochrome *c*.. The red lines across the temperature table and insert highlight the drastic drop in Tm values of AaCPR1 across the tested eight cytochrome *c* concentrations. **B.** AaCPR2 gradually decreased its melting temperature with increased cytochrome *c* concentrations. No initial transition temperature was recorded at any of the cytochrome *c* concentrations. The sole melting peak lost energetic dominance as the ligand concentration increased.

Increasing concentrations of cytochrome *c* gradually decreased the second melting temperatures of the two enzymes: from 61.3°C to 46.5 °C for AaCPR1 and from 54.2 °C 47.7°C for AaCPR2 (Fig. 9A-B). AaCPR1 and AaCPR2 also exhibited different responses in the first melting temperature to the increase of cytochrome *c*. When the concentration of cytochrome *c* was increased to 10μM, the first melting temperature of AaCPR1 became unstable and was essentially lost for all subsequent concentrations of cytochrome *c* (Fig. 9A). Conversely, the melting temperature of AaCPR2 remained monophasic throughout all concentrations, gradually losing energetic dominance, particularly at 100μM of cytochrome *c* (Fig. 9B). These results reveal inhibitory effects of high cytochrome *c* concentrations on the unfolding capacity for both enzymes.

### Structural Simulation of AaCPR1 and AaCPR2 with FAD, NADP(H), and FMN

AaCPR1 and AaCPR2 structures were predicted using ColabFold (CF) (Mirdita et al., 2022) and aligned to the crystal structures of ATR2 (<u>5GXU</u>) and SbCPR2b (<u>7SV0</u>) from the protein data bank for proper positioning of the FAD, FMN and NADP^+^ ligands (Fig. 10). No NADPH fitting exists among the plant crystal structures. This is due to the transient nature of NADPH interaction at the binding site (Niu et al., 2017; Whitelaw et al., 2015). Consequently, the binding of NADP^+^ reflects AaCPR1 and AaCPR2 in their semi-reduced form. Nonetheless, the SbCPR2b structure serves as a good proxy for NADPH binding. The distance in the predicted models between the N5 positions of FAD and FMN were 12.689 Å and 12.775 Å for AaCPR1 and AaCPR2. This is nearly half the distance reported for crystallized ATR2 22.9 Å from N5 to N5, demonstrating that ColabFold predicts the CPRs in a more closed state. This is supported by other crystal structures such as rat CPR (<u>1JA0</u>) in its closed conformation with a 13 Å N5 to N5 distance. Furthermore, Cryo-EM data on rCPR shows that different particle sets captured the enzyme across several closed conformations (Lepesheva et al., 2026). Therefore, previous crystal structures and ColabFold depict reasonable, yet static representations of CPRs and the literature reported distances between the FAD and FMN co-factors still serve as appropriate references.

**Figure 10.**
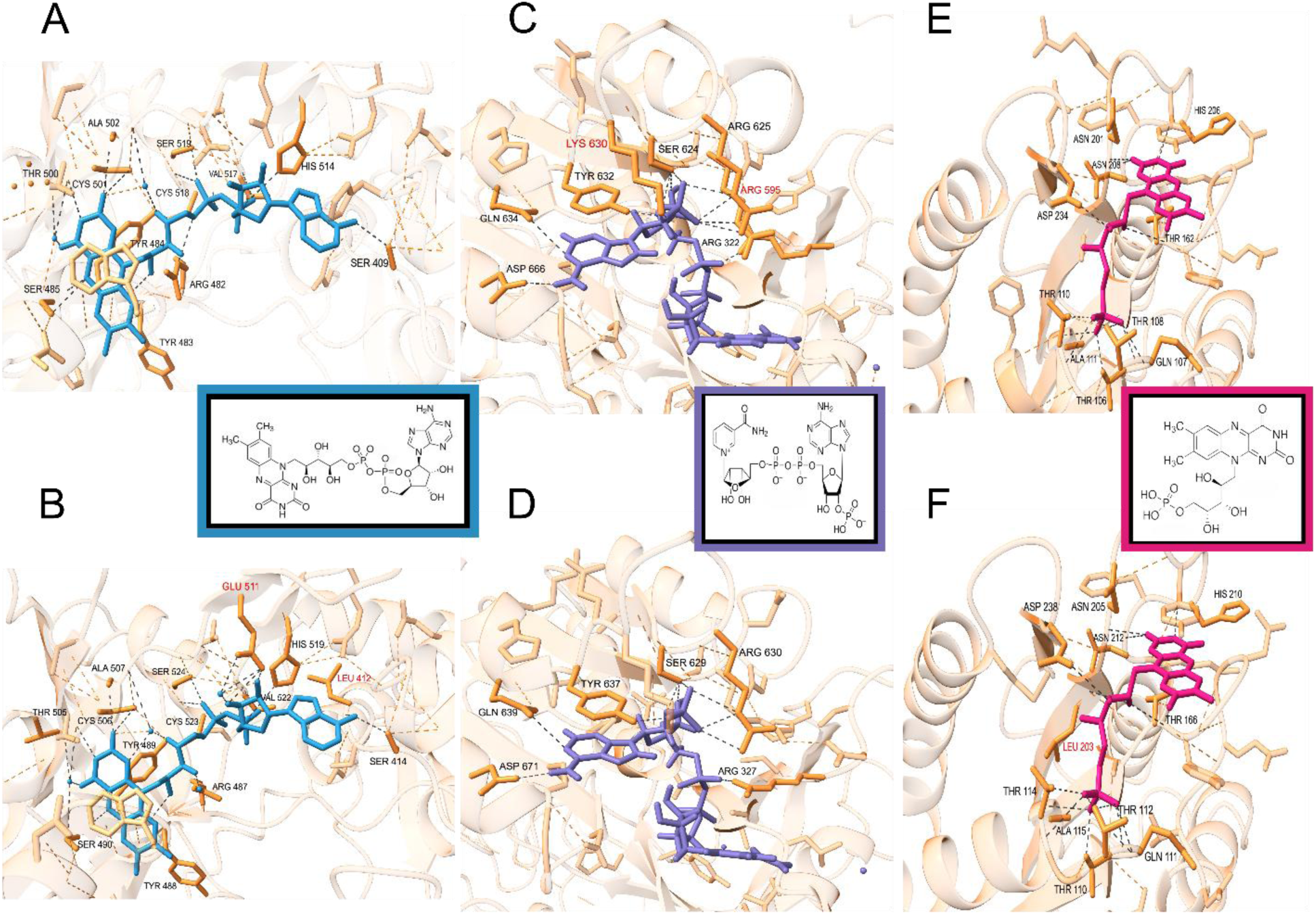
**Docking simulation showing interactions of FAD, NADP^+^ and FMN ligands with AaCPR1 and AaCPR2**. ColabFold predicted models were aligned to ATR2 (5GXU) and SbCPR2b (7SV0) for ligand positioning. Residues within 5 Å of the ligands are shown. Those amino acid residues forming hydrogen bonds with the co-factors are labeled in black. Directly interacting hydrogen bonds are highlighted with dark grey and all other bonds in the pocket are highlighted with orange. Inserts show the corresponding molecular structures of the fitted co-factors. **A-B.** FAD within the FAD binding region of AaCPR1 (A) and AaCPR2 (B). LEU412 and GLU511 are highlighted with red colour to show that they form bonds with FAD in AaCPR2 but not in AaCPR1 model. **C-D.** NADP^+^ within the NADP binding region of AaCPR1 (C) and of AaCPR2 (D). The hydrogen bonds of ARG595 and LYS630 highlighted with red colour in AaCPR1 were not detected in the AaCPR2 model. **E-F.** FMN within the FMN binding domain of AaCPR1 (E) and AaCPR2 (F). Hydrogen bonding of LEU203 formed in AaCPR2 highlighted with red colour was not detected in AaCPR1.

When the CF-predicted AaCPR FMN domains are aligned to the ATR2 FMN domain, the resulting distance between FAD and FMN increases, and key interfacing residues present on the loops between structural elements are exposed. This includes GLN112, TYR168, ASP175, ASP666, and LYS 668 (ATR2 numbering) (Niu et al., 2017) (Fig. S5). These loops represent the region of the FMN binding domain exposed to solvent (Huang et al., 2015). The CF models of AaCPR1 and 2 also share highly conserved residues within the FAD binding pocket, forming hydrogen bonds (Fig. 10A-B), which are nearly identical to residues present in ATR2 (Niu et al., 2017). There are only two differences between ATR2 and the AaCPRs at this binding site.

Firstly, AaCPR1 and AaCPR2 include an additional cysteine bond at O2 of the FAD isoalloxazine ring. Secondly, there is a switch from phenylalanine at position F490 in ATR2 to a tyrosine at position Y483 in AaCPR1 and Y488 in AaCPR2. This residue forms a conserved bond with the hydroxy group of FAD’s riboflavin opposite tyrosine Y491 in ATR2 and Y484/Y489 in AaCPR1 and AaCPR2. Additionally, the predicted two models show that the FAD binding pocket of AaCPR2 possesses two additional hydrogen bonds formed at LEU412 and GLU511, while AaCPR1 does not (Fig. 10B). Of these two bonds, LEU412 hydrogen bonding to the amine of FAD’s adenine is present in ATR2 (L414) (Niu et al., 2017), while GLU511 is absent in ATR2.

According to the structure of ATR2, the features of the NADP pocket of both AaCPR1 and AaCPR2 were compared to reveal their differences and similarities. The resulting models showed that THR570, which forms a bond at the second phosphate of the ribose-adenine ring in ATR2, was not present in either of the CF predicted AaCPR1 or AaCPR2 models. AaCPR1 shared the greatest number of residues with ATR2, including LYS630 (LYS637 ATR2), which was present but not hydrogen bonded to NADP^+^ in AaCPR2 (Fig 10C-D). Additionally, NADP^+^ in AaCPR1 established bonds with ARG595, a residue bonded in neither the ATR2 crystal, nor the AaCPR2 prediction (Fig. 10C). Apart from these differences, the NADP pocket of AaCPR2 shared all other hydrogen bonding residues with ATR2 and AaCPR1 (Fig. 10D). Notable non-hydrogen bonding amino acid motifs in the AaCPRs includes the “Asp-loop” (Hubbard et al., 2001; Zhang et al., 2022): ^658^GDAKGM^663^ for AaCPR1 and ^662^GDAKGM^667^ for AaCPR2. The signature ASP659/663 residue is positioned near the positively charged nicotinamide ring, as reported in the SbCPR2b crystal structure (Zhang et al., 2022). This loop has been proposed as a key factor in the binding of NADPH and release of NADP^+^ (Zhang et al., 2022).

The FMN pocket likewise maps all the same residues between CF-predicted AaCPR1 and AaCPR2 structures (Fig. 10E-F). The GLN107 (AaCPR1)/GLN111 (AaCPR2) residue, which is implicated in FAD-FMN electron transfer (Niu et al., 2017), also forms hydrogen bonds with FMN in the enzyme binding pockets. Rather than a glycine (G114), as observed in ATR2, the models uncovered that AaCPR1 and AaCPR2 possess an alanine, A111 and A115, respectively, which interact with the terminal phosphate of FMN. The CF models also yielded fewer interactions between residues and the FMN isoalloxazine ring as reported in ATR2 (Niu et al., 2017); however, the interacting residues in ATR2 are also conserved in the binding pocket of the AaCPRs, such as the surrounding tyrosines and a glycine on one of the domain loops. Lastly, the CF AaCPR2 differs between AaCPR1 and ATR2 by forming a hydrogen bond between L203 and the second hydroxy group of FMN (Fig. 10F).

The models of AaCPR1 and AaCPR2 were further analyzed to characterize the NADPH binding region and connecting domain interactions that flank each side of the FMN domain and contain essential interfacing residues. The resulting data showed that the residues in these regions for AaCPR1, AaCPR2, and ATR2 are identical. Of particular interest is the conservation of the plant specific R708-E186 salt bridge between the NADPH and FMN binding domains reported in ATR2 (Niu et al., 2017). The mutation of R708 was reported to completely abolish interactions with cytochrome *c*, determining its essential role for electron transfer from FAD to FMN in plant CPRs (Niue et al., 207). The models show that this bridge is present in the AaCPRs as R706-E185 (AaCPR1) and R701-E181 (AaCPR2). These data indicate that both AaCPR1 and AaCPR2 are likely to use this bridge to interact with cytochrome *c* to perform their reduction activities. Altogether, these results show differences between ATR2 and the AaCPRs and between the isoforms themselves.

## Discussion

### AaCPR1 and AaCPR2 Group Into the Same Clade

The phylogenetic tree revealed that AaCPR1 and AaCPR2 group into the same clade. Given the canonical understanding of the functional differentiation between CPR1 and CPR2, the co-existence of the AaCPR paralogs in the CPR2 implies that they share similar functions (e.g., inducible expression). Despite their high phylogenetic similarities, a wider phylogeny with additional plant CPR sequences reveals that AaCPR1 groups more closely to CPR2a from *Helianthus tuberosus* (Jerusalem artichoke) and AaCPR2 more closely groups with CPR2 from *Stevia rebaudiana* (stevia/candyleaf) (S6), both of which are involved in the formation of sesquiterpene lactones (Borgo et al., 2023; Yuan & Yang, 2017). This phylogenetic feature not only suggests similar functions of the two paralogs, but also shows their potential divergence, supporting the importance of an in-depth investigation into their metabolic differences (e.g., comparing activities of the AaCPRs paralogs in association with CYPs from artichoke and stevia). Experiments against UV-B and UV-C light among *A. annua* plants showed that high radiation triggers *AaCPR1* expression in association with other artemisinin genes such as amorpha 4,11-diene synthase (ADS), and CYP71AV1 (Rai et al., 2011), and expression data has demonstrated differential expression between the *AaCPRs* (Ma et al., 2015). The expression level of *AaCPR2*, in particular, was highest in flowers where artemisinin content was reported to be the most abundant (Ma et al., 2015). This suggests that, despite their high degree of relatedness, AaCPR1 and AaCPR2 are metabolically divergent *in planta*. Therefore, immediate next investigations are anticipated to elucidate their contributions to artemisinin biosynthesis and redefine classification of AaCPR1 in the plant CPR2 clade.

### Residue Variability at the N-terminus, Hinge, and Connecting Regions of AaCPRs May Impact Their Protein/Substrate Interactions

At the amino acid sequence, the two paralogs share around 70.8% identity. A major source of their variation is localised at the N-terminus (Fig. S1), which is associated with membrane integration and facilitation of P450 interactions for peak electron transfer activity (Park et al., 2013; Strohmaier et al., 2019). Park et al. (2013) suggest that amino acid differences in this region could vary the electrostatic potentials of the enzymes, facilitating different P450 interactions. While the two paralogs share conserved sequences of FMN, FAD, and NAD binding motifs, differences exist in other regions. Like other known plant CPRs, AaCPR1 and AaCPR2 vary in amino acid identity at their flexible hinge. The alignment undertaken in this work revealed that the hinges of the two enzymes differ by two amino acids (Fig. S1). First, T262 in AaCPR1 is substituted by M266 in AaCPR2, and second, A273 is substituted by L277 at the corresponding position (Fig. S1). These substitutions replace hydrophilic threonine with a hydrophobic methionine and a short side-chain alanine with a bulkier leucine. Since the hinge facilitates rotation of the FMN domain toward and away from the FAD during electron transfer, even subtle differences within this region can affect the reduction capability of CPRs (Campelo et al., 2017; Im & Waskell, 2011; Laursen et al., 2011; Niu et al., 2017; Wang et al., 1997). For instance, in human CPR, a hinge residue mutation from alanine to isoleucine resulted in lower cytochrome *c* reduction rates when the protein was in its membrane bound form (Campelo et al., 2017). This supports the understanding that side-chain size impacts hinge flexibility and rigidity, thereby differentially affecting electron transfer (Campelo et al., 2017; Kakraba et al., 2026).

The connecting domain is a region that introduces the greatest amount of residue variability between AaCPR1 and AaCPR2 (Fig. S1). This domain has also been implicated in facilitating FAD and FMN interactions and subsequent FMN domain movement (Aigrain et al., 2011; Laursen et al., 2011; Wang et al., 1997). It characteristically shows the lowest sequence homology between CPRs (typically less than 30%), which is reflected through differences in their secondary structure (Murataliev et al., 2004; Waskell & Kim, 2015). The largest uninterrupted segment of variable residues (after the N-terminus) between AaCPR1 and AaCPR2, is a loop that shortly follows the hinge: AaCPR1’s ^285^ETYDQDQ-L^292^, and AaCPR2’s ^289^DSSAEDHSH^297^ (Fig. S7). This segment adopts neither α-helical nor β-sheet secondary structure, and the residue composition of AaCPR1 and AaCPR2 is consistent with dynamic, flexible, solvent-exposed loop regions (Mitusińska et al., 2020). (Fig. S7). Little is known about the functional contributions of this loop because neither the ATR2 nor the SbCPR2b crystal structures have been able to co-crystallize with it (Niu et al., 2017; Zhang et al., 2022). This is largely attributed to the flexible nature of these regions as they are especially difficult to resolve with X-ray crystallography and cryogenic electron microscopy (Cryo-EM) (Askenasy et al., 2018; Huang et al., 2013; Mitusińska et al., 2020). When full-length rat CPR was submitted to Cryo-EM, its N-terminal anchor lacked sufficient density, which was attributed to its high mobility in solution (Lepesheva et al., 2025). This introduces the possibility that the ^285^ETYDQDQ-L^292^ and ^289^DSSAEDHSH^29^ loops from the AaCPRS, which lead into the connecting domain, may also possess flexible capacities like the hinge and N-terminal anchor (Aigrain et al., 2012; Lepesheva et al., 2026; Wang et al., 1997). The lack of conservation in this region suggests that this loop, much like what has been observed for the N-terminal anchor, is inconsequential for electron transfer (Murataliev et al., 2004). However, just as the anchor has been identified as a requirement for most CYP interactions (Djordjevic et al., 1995; Laursen et al., 2011; Miyamoto et al., 2015; Murataliev et al., 2004; Strohmaier et al., 2019; Wang et al., 1997), and the hinge region is emerging as an essential component for establishing CYP preferences, it is possible that this loop may contribute to differential CPR conformational dynamics or selection. The 285-292 segment of AaCPR1 contains leucine and tyrosine which have more hydrophobic properties, whereas the corresponding 289-297 segment of AaCPR2 has three polar serine residues that are hydrophilic. While both sequences are negatively charged, the larger size of TYR287 and L292 in AaCPR1 may constrain loop movement more than that of the serine-rich AaCPR2 sequence that holds residues with smaller side chains. Altogether, the differences herein offer potential explanations for the different biochemical and biophysical properties observed between the two enzymes.

### AaCPR1 and AaCPR2 Both Form a Charge-Transfer Complex and Differ in Their Kinetic Traits

AaCPR1 and AaCPR2 behaved similarly during spectral analysis (e.g., shared asymmetric peaks from 450-480 nm), indicating a similar flavin content for both (Fig. 5). Neither could be fully reduced by NADPH. This is indicative of reoxidation events occurring from the formation of NADP^+^ species as reactants and products become equivalent, creating a mixture of CPRs in their semi-reduced and reduced states (Vigil et al., 2021; Whitelaw et al., 2015). Consequently, the absorption capacity of the AaCPRs was expanded from 550 nm to 700 nm, which is associated with the formation of the NADP^+^-FADH_2_ charge transfer complex (Whitelaw et al., 2015; Zhang et al., 2022). This has been widely documented in stopped-flow experiments (Simtchouk et al., 2013; Whitelaw et al., 2015). Once established, NADP^+^-FADH_2_ prevents the reduction of FMN—this slows interflavin transfer among plant CPRs. Subsequent ET steps are reliant on the displacement of NADP^+^ by a conserved tryptophan that stabilises the formation of the FADH semiquinone after the initial one-electron transfer to FMN (Whitelaw et al., 2015). Both AaCPR1 and AaCPR2 have this residue at their C-terminus (W704 and W709), which is consistent across all plants (Whitelaw et al., 2015; Zhang et al., 2022). The loss of NADP^+^-FADH_2_ in plant CPRs was reported to take 20-50 seconds and is thus dictated as a rate-determining step for cytochrome *c* reduction (Whitelaw et al., 2015). Thus, accelerating the speed at which NADP^+^ is displaced is a tenable kinetic target for future plant CPR engineering strategies. Consequently, the kinetic differences observed between AaCPR1 and 2 could be attributed to differences in the way the respective enzymes form and disassociate from the charge-transfer complex. AaCPR2 demonstrated greater acceleration at saturating conditions with a higher V_max_ and k_cat_ when assayed with cytochrome *c* and NADPH, while AaCPR1 possessed a higher enzyme efficiency, based on its lower K_m_ and higher k_cat_/K_m_ values (Table 1). Thus, the differing substrate and co-enzyme saturation requirements observed between AaCPR1 and AaCPR2 may reflect differences in the formation, stabilisation, or dissociation dynamics of the NADP^+^-FADH_2_ charge-transfer complex during catalytic cycling. Consequently, the distinct kinetic requirements of AaCPR1 and AaCPR2 support the hypothesis that these enzymes are functionally divergent.

### Characteristics of AaCPR1 and AaCPR2 Thermostability Based on Differential Scanning Fluorimetry

Differential Scanning Fluorimetry (DSF)/Thermofluor characterised the thermostability and unfolding transitions of AaCPR1 and AaCPR2 by two aspects. The first is that the two proteins possess different baseline melting/unfolding temperatures when unbound. The other is that their physical features are impacted by the type and concentration of the ligands present. (Fig. 7-9). The melting temperatures of AaCPR1 were consistently higher than those of AaCPR2 under all tested conditions, indicating greater overall stability. Furthermore, AaCPR1 had a persistent biphasic (double peak) melting pattern, particularly at low NADPH and cytochrome *c* concentrations. This biphasic behaviour has been reported previously for flavin containing enzymes from other species, suggesting that enzyme stability is a characteristic difference among flavodoxin/diflavin paralogues (Lawson et al., 2004). The following discussion herein adopts the presumption that CPR stability, as represented in the Thermofluor data, may be reflective of enzyme conformational changes, such that: lower thermal stability is associated with an extended/open conformation and higher thermal stability is associated with a closed conformation (Huang et al., 2013; Iijima et al., 2019). Furthermore, within the context of electron flux, open conformations are thought to correspond to cytochrome c-reactive states, whereas closed conformations are thought to correspond to comparatively unreactive states (Haque et al., 2014).

An interesting biophysical phenomenon regarding the biphasic nature of AaCPR1 and AaCPR2 was that both were ligand- and dose-dependent (Fig. 7-9). For example, NADH, when applied under any condition tested with or without cytochrome *c*, caused both enzymes to transition to a clear monophasic melting pattern (Fig. 7). Furthermore, the application of NADH abruptly reduced the melting temperatures of both enzymes. These features coincide with their poor kinetic performance in the steady state assays with NADH, which is indicative of weak binding (Table 1). This suggests that weak co-enzyme binding correlates to lower thermal protein stability. Based on a previous report from Huang et al. (2013), NAD^+^ is capable of reducing human CPR to CPR^2e−^; however, the weak binding of NAD^+^ yields a co-enzyme free structure. In small angle x-ray scattering (SAXS) experiments with hCPR and NADH, the protein structures assumed a less compact conformation. Thus, the absence of co-enzyme binding is attributed to a less stable/open structure, while the presence of co-enzyme binding results in a more stable/compact structure (Huang et al., 2013). By contrast, NADPH increased the major melting temperatures for both AaCPR1 and AaCPR2 (Fig. 7-8). This increase is congruent with improved enzyme thermal stability via formation of the charge transfer complex, wherein NADP^+^ remains bound to the NAD binding site (as reported previously) (Simtchouk et al., 2013; Whitelaw et al., 2015). In AaCPR1, higher NADPH concentrations (≥ 30μM) led to the loss of biphasic melting (Fig. 8). The biphasic melting changes of AaCPR2 were less prominent (Fig. 8). Though, two peaks were observed at ≥ 2μM of NADPH, their nearly equal depths/energetic dominance challenges the designation of two discrete melting transitions. One consideration is that different peaks are associated with different redox species (e.g., NADP^+^-FADH_2_, flavin semiquinones, and flavin hydroquinones), and by extension, different conformational CPR states (Huang et al., 2013). Thus, establishment of a persistent redox specie/protein conformation could account for the disappearance of biphasic melting behaviour in the two enzymes. The singular melting transition primarily consistent with AaCPR2 may then indicate its propensity to dominate one redox/conformational state over all others. The specific type of redox state/conformation that results in this thermal response, however, remains open for investigation.

Another unique thermo-structural difference between the two proteins was their response at 100-200μM NADPH (Fig. 8). Higher concentrations of NADPH increased the second melting temperature of AaCPR1 but eliminated the first melting temperature. Conversely, at 100-200μM NADPH, AaCPR2 resumed a biphasic melting pattern, where the first melting transition was equally or more dominant than the second melting peak (Fig. 8, ReplicateCurves_DSF.pdf).

Although the nature of these changes requires further characterisation, one hypothesis is that the lower thermal stability of AaCPR2 was associated with weaker affinity/binding of NADPH/NADP^+^ at these concentrations. It is possible that this forms a milieu of co-enzyme bound (closed) and non-co-enzyme bound (extended) conformations (Huang et al., 2013); hence, the two peaks. Thus, the thermostability differences between AaCPR1 and AaCPR2 under NADPH conditions may be reflective of their divergent affinities for NADPH and their ability to bind and maintain binding with the co-enzyme. This demonstrates a diverse dose dependent stability for AaCPR2 against NADPH and emphasizes its more reactive behaviour to ligand changes. The diverging behaviour of AaCPR1 and AaCPR2 may also result from differential preferences for arrangements of the charge-transfer complex. Zhang et al. (2022) reported that NADP^+^ situates within the binding pocket differently depending on the redox state of SbCPR2b. When SbCPR2b was fully oxidised, the nicotinamide moiety of NADP^+^ was localized in the binding pocket near E508 and R323 in a more extended (syn) conformation, which increased the distance between the moiety and the isoalloxazine ring. By contrast, when SbCPR2b was in its naturally oxidized form, the nicotinamide moiety shifted closer to the isoalloxazine ring in an *anti*-conformation near D661. These positional movements result in not only the conformational alterations of plant CPRs but also affect their hydride transfer properties (Murataliev et al., 2004; Zhang et al., 2022). Taken together, the dose-dependent responses of both AaCPR1 and AaCPR2 provide useful information for elucidating their roles in artemisinin biosynthesis in the future.

The behaviour of the AaCPRs in their oxidised form (i.e., without reduction by NADPH) was studied with the addition of cytochrome *c* by itself (Fig. 9). Without NADPH to reduce AaCPR, the assumption was that the protein would not vary in thermal stability and thus resemble its oxidised ligand-free state. However, increasing concentrations of cytochrome *c* decreased the melting temperatures of both AaCPR1 and AaCPR2, and consequently, their thermal stability (Fig. 9). This finding was slightly unexpected as structural and molecular dynamics simulations on rat CPR have shown that it predominantly assumes a closed (unreactive) state when oxidised and thus ought to align with a more thermally stable state (Haque et al., 2014; Huang et al., 2013; Iijima et al., 2019). An important caveat to these structural investigations, however, was that they did not include structural inference of oxidised CPR when exposed only to cytochrome *c* (Haque et al., 2014; Huang et al., 2013). A negative control performed during the AaCPR kinetic experiments showed that co-enzyme free AaCPR and cytochrome *c* were non-reactive. Therefore, the thermal results in this case are not the result of spontaneous reduction of cytochrome *c*. This suggests that there may be a non-redox dependent mechanism that caused the reduced thermal stability (protein opening/extension) of the oxidised AaCPRs in the presence of cytochrome *c*.

Supplemental experimentation with fixed cytochrome *c* (20μM) and increasing NADPH concentrations resulted in less dynamic temperature shifts (Fig. S3, ReplicateCurves_DSF.pdf). These conditions resulted in closely overlapping melting peaks for both enzymes (Fig. S3A).

This implies that 20μM cytochrome *c* largely influenced AaCPR melting behaviour regardless of how much NADPH was present. In AaCPR1 this was reflected by the predominating initial melt curve at ∼36.5°C, which indicates lower thermal stability (Fig. S3A). This may reflect a tendency for AaCPR1 to assume a more dominant open and reactive cytochrome *c* conformation under these conditions. Interestingly, AaCPR2 maintained a more predominant biphasic melting curve (Fig. S3B). This could be the result of a persistent equilibrium forming between AaCPR open and closed states. This corroborates the observation among human diflavin enzymes (CPR included), that a mixture of open and closed conformations forms when the enzymes are fully reduced in the presence of cytochrome *c* (Haque et al., 2014). Whether these observations correlate to the respective substrate affinities between the two enzymes (i.e., a lower K_m_ value for AaCPR1 and higher K_m_ value for AaCPR2) will require further experimentation.

When NADPH remained fixed (30μM) and cytochrome *c* was increased, the melting curves were more dynamic (Fig. S3B). An increase of the second melting transition was observed in both AaCPRs when cytochrome *c* was ≤ 2μM (Fig. S3B). More notably, was the near total loss of fluorescence/observed unfolding among the enzymes when cytochrome *c* was ≥ 50μM (Fig. S3B). This could be the result of a co-enzyme and substrate induced stabilisation that is resistant to the Thermofluor temperature changes/undetectable by the assay (Pinz et al., 2022).

Altogether, the differences in melting behaviour across Thermofluor conditions seem to suggest that the conformational dynamics of AaCPRs are substrate and concentration dependent.

### Influence of Molecular Bond Differences Between AaCPR1 and AaCPR2 on Conformational Change

The source of the biophysical differences between AaCPR1 and AaCPR2 is challenging to pinpoint on a structural level. Primarily, the ligand pockets modeled by ColabFold in Figure 10 show minimal differences between the non-covalent bonds of the two enzymes in either the FAD, NADP, or FMN bound models. In fact, many of the prominent residues in these sites are also featured in ATR2 from Arabidopsis (Niu et al., 2017). This affirms that highly conserved binding sites for co-enzymes and co-factors are essential for CPR function but offers little insight into the different biophysical properties of the enzymes. From what is known in the literature, different CPR redox states affect positioning of ligands—sorghum SbCPR2b was only co-crystallized with NADP^+^ when the enzyme was in its oxidised state and attempts to capture the exact positioning of NADP^+^ when rat CPR was fully reduced were unsuccessful (Xia et al., 2018; Zhang et al., 2022). Indeed, no NADPH-bound CPR crystal structure exists (Xia et al., 2018; Zhang et al., 2022). This suggests, thereby, that molecular orientations of NADPH and possibly other redox states of the FAD and FMN co-factors may not be captured either by known crystal or ColabFold predicted models. Thus, it is possible that the unknown orientations of NADPH form other hydrogen bonds with different residues in the AaCPR proteins, resulting in a wider breadth of, or different types, of hydrogen bonds which are not detected by the available models. This, in turn, could explain the different biophysical responses of AaCPR1 and AaCPR2 to NADPH.

Intra-domain movements have also been observed in rat FMN domains, wherein a fully reduced (four-electron) CPR resulted in a rotation of the TYR140 sidechain that sits below the isoalloxazine ring (Xia et al., 2018). While this rotation effectively breaks hydrogen bonds at the phosphate group of FMN, the overall domain movements result in tighter interdomain interactions between the FAD and FMN groups as the broken bond enables movement of the GLY141 loop (Xia et al., 2018). This loop is an essential interfacing FAD region in ATR2 (Niu et al., 2017) and was confirmed in the AaCPR proteins through homology modeling, alongside residues TYR163 and TYR167 in AaCPR1 and AaCPR2 (Fig. S5). These shared residues suggest that this FMN associated domain movement is conserved across species. Additionally, NADPH reduced rat CPR results in retraction of the previously described Asp-loop to form additional hydrogen bonds with the FMN ring (Xia et al., 2018). Such mechanisms explain the improved thermostability observed in AaCPR1 and AaCPR2 when NADPH was added to the Thermofluor assay (Fig. 7). Whether the co-occurrence of increasing NADPH concentrations and enzyme stability is a result of increasing reduction potential in the enzymes, or a more physical based mechanism due to ligand abundance requires further investigation. These documented redox-induced domain and ligand changes in CPRs establish the likelihood of other intradomain motions that have yet to be characterised, which may be reflected in the varying Thermofluor melting curves. The slight differences observed in the hydrogen bonding of residues to the CF-predicted models of AaCPR1 and AaCPR2 also reflect how minor ligand translations and rotations may differentially impact the bio-structural organization of the enzymatic domains.

## Conclusion

Altogether, different redox states, the positional changes of transient ligands, and residue differences and domain motions could help to explain the differences observed between AaCPR1 and AaCPR2 during the enzyme kinetic and Thermofluor assays. These characteristics could consequently bestow a functional bias between the two CPRs against different interacting CYPs. This is supported by studies on plant CPR paralogues and closely related plant CPRs that demonstrate diverging catalytic activities and even functional exclusivity toward a CYP substrate (Ponnamperuma & Croteau, 1996; Su et al., 2017). A study by Jensen et al. (2021) proposed the concept of biased metabolism wherein CPRs attenuate their synergistic and inhibitory functions with CYPs depending on what small molecules bind to them. This was attributed to changes in the conformational equilibrium of the CPR forms—one shape persisted in greater abundance over the others, which resulted in a preferential partnership with a particular CYP (Jensen et al., 2021). Transitions between CPR conformational states occur on the magnitude of nanoseconds to milliseconds, so it is not possible for the Thermofluor assay described here to resolve all the possible states (Jensen et al., 2021). However, based on SAXS data (Huang et al., 2013), discrete CPR populations in their closed and open conformations, existing in equilibrium, could explain the biphasic melting curves observed in AaCPR1 and AaCPR2, particularly in the combinatorial NADPH-cytochrome *c* assays. Thus, the biophysical differences between AaCPR1 and AaCPR2 communicate a deeper functional reality—that these enzymes prioritize different CPR conformations to facilitate divergent metabolic outcomes with conformation driven differential CYP selections. This idea relates to emerging theories that identify CPR as a more active participant in CYP metabolism and qualifies plant CPR paralogues as metabolically significant participants in diversifying plant metabolism. Uncovering these differences will have a positive impact on the further elucidation of artemisinin biosynthesis and the roles that CPR paralogues have in the production of other terpenoids. In future experiments, amino acid alterations in the ^285^ETYDQDQ-L^292^ and ^289^DSSAEDHSH^29^ loops will be evaluated to test their functions in redox reactions for the biosynthesis of artemisinin.

## Experimental Procedures

### Materials

Oligomeric primers for gene cloning were synthesized from either MilliporeSigma (Burlington, MA) or ETON Biosciences (Durham, NC). Chemical agents were purchased from different companies, including ampicillin and kanamycin from Sigma-Aldrich (St. Louis, MO), IPTG and Ni-NTA from Goldbio (St. Louis, MO), lysozyme from VWR International (Radnor, PA), Tris, NaCl, and imidazole from Thermo Fisher (Durham, NC), EDTA, DTT, bovine serum albumin, NADPH, NADH, and cytochrome c from equine heart by Sigma-Aldrich (St. Louis, MO). Coomassie Blue R-250, protein ladder, and SDS reagents from BioRad (Hercules, CA). Amylose resin and maltose were purchased from NEB Biolabs (Ipswich, MA). SYPRO™ Orange protein gel stain was purchased from Thermo Fisher Scientific (Eugene, OR).

### Phylogenetic analysis

Thirty-three CPR amino acid sequences from various plant species were identified from the literature and NCBI database (Supplementary Table 1). Sequences were aligned with ClustalX (Thompson et al., 1994) and used to generate a maximum likelihood phylogenetic tree with 1000 bootstrap replications in [Mega11](https://www.megasoftware.net/) (Tamura et al., 2021). The generated tree file was uploaded into [ITOL](https://itol.embl.de/) (Interactive Tree of Life) for further annotation.

### Cloning of truncated AaCPR1 and AaCPR2

RNA was extracted from *Artemisia annua* inflorescences using the <u>RNeasy extraction kit</u> (Qiagen, USA) and used to synthesize the first strand cDNAs with Five prime Rapid Amplification of cDNA Ends <u>(RACE) SMARTer kit</u> (Clontech, USA). The resulting cDNAs were used as template for polymeric chain reactions (PCR) to amplify the open reading frames (ORF) of *AaCPR1* and *AaCPR2,* based on the complete sequences for AaCPR1 (<u>EF197890.1</u>) and AaCPR2 reported by Ma et al (2015). The complete ORFs of each gene were used as templates for a second PCR reaction to truncate the first 198 bp (encoding 66 amino acids) and 201 bp (encoding 67 amino acids) of *AaCPR1* and *AaCPR2*. The primers used for these cloning steps are listed in Table S1. AaCPR2 also included a TEV sequence prior to the truncated ORF. The respective ORFs were then cloned using the <u>Gateway Cloning technology</u> from ThermoFisher Scientfic (Waltham, MA) into the <u>pDEST17</u> vector for AaCPR1 and the <u>pDEST-HisMBP</u> for an MBP tagged AaCPR2.

### Recombinant enzyme expression and purification

AaCPR1 and AaCPR2 plasmids were transformed into competent *E.coli* <u>Rosetta BL21(pLysS)</u> <u>cells</u> from ThermoFisher and screened on agar-solidified LB medium supplemented with 100μg/mL ampicillin. Positive colonies were inoculated into 25mL of LB broth containing ampicillin (100μg/mL) and grown to a density of approximately O.D._600 =_1.0 overnight in a 37°C shaking incubator. 1-3 mL of this starter culture was used for the inoculation of four 500 mL volumes of TB media and 100μg/mL ampicillin. These flasks were grown for 6-7 hrs at 37°C in a shaking incubator until the O.D._600_ reached 0.8, after which, 0.1 mM IPTG was added to each flask. Flasks were subsequently moved to a 14°C shaking incubator and left to induce the proteins for 44hrs. Cells were then harvested using an <u>Avanti JXN-26 centrifuge</u> (Beckman Coulter, USA) by spinning them down at 8000rpm for 30min at 4°C . Cell pellets were washed in a suspension of 25 mL dH_2_O before being centrifuged under the same conditions and placed in 50mL falcon tubes for storage in a −80°C freezer until protein extraction and purification.

Frozen pellets amounting to ∼10-15g were resuspended in 200 mL of extraction buffer consisting of 50 mM Tris-HCl pH 7.5, 200 μg/L of lysozyme, 1mM phenylmethylsulfonyl fluoride (PMSF), and 1mM benzamidine. This mixture was kept on ice for 30 min before lysing with a <u>120 Sonic Dismembrator</u> (Fisher Scientific, USA) set to 40% AMPS for 5min 20s on/10s off.

This setting was repeated until the cell mixture was fully homogenous. After centrifugation to remove all cell debris, the supernatant was used for protein purification.

### Purification of AaCPR1

The purification of AaCPR1 was performed as described by Simtchouk et al. (2013) with minor revisions. In brief, the crude preparation of AaCPR1 cell lysate was combined with 150mL of column buffer (50mM Tris-HCl pH 7.5, 500mM NaCl, 20mM imidazole) and loaded onto a 20mL Ni-NTA column equilibrated with 10 volumes of column buffer. The column was washed with 1400mL of column buffer and protein was eluted with 50mM Tris-HCl pH 7.5 and 300mM imidazole. Yellow elution fractions were pooled, confirmed via SDS-Page and concentrated with an <u>Amicon ® Ultra Centrifugal Filter (30kDa MWCO)</u> (Sigma-Aldrich, USA). The concentrated samples were submitted to additional purification using an <u>ÄKTA pure™ chromatography</u> <u>system</u> (Cytiva, Sweden) with a 1mL <u>HiTrap IMAC high performance column</u> (Cytiva, Sweden).

Subsequent concentration and desalting of protein was achieved by passing the samples through the 30kDa filters and removing imidazole from elution buffer with 50mM Tris-HCl pH 7.5 up to three times. The purity of the sample was assessed via SDS-PAGE and protein concentration was determined using the Bradford method before being flash frozen in liquid nitrogen and stored in - 80°C.

### Purification of AaCPR2

AaCPR2 cell lysate was added to 200mL of column buffer (50mM Tris-HCl pH 7.5) and loaded onto an equilibrated 20mL Ni-NTA column. The column was washed with 1800mL of column buffer and then 2L of wash buffer (50mM Tris-HCl pH 7.5, 200mM NaCl, 20 mM imidazole). Protein was eluted with 50mM Tris-HCl pH 7.5 and 300mM imidazole. Yellow elution fractions were pooled and added to 50mL of a TEV protease buffer (50mM Tris-HCl pH 7.5, 0.5mM EDTA, 1mM DTT). <u>IceTEV protease</u> from Sigma-Aldrich was added at 1:40 TEV to target protein ratio to this solution and then left overnight at 4°C. The digestion was then prepared in 10mL aliquots (40mg protein) and then loaded onto a 5 mL amylose-MBP affinity column equilibrated with 10 volumes of column buffer (20mM Tris HCl pH 7.5, 200mM NaCl, 1mM EDTA) to remove the cleaved MBP tag and other non-target proteins. The MBP-free column flow-through was concentrated and desalted with 30kDa filters. The purity of the sample was assessed via SDS-PAGE and protein concentration was determined using the Bradford method before being flash frozen in liquid nitrogen and stored in −80°C.

### Absorbance Spectroscopy and Reduction of CPR with NADPH

The spectral properties of the two CPR enzymes were characterized anaerobically in a <u>Coy Lab</u> <u>Vinyl Chamber</u> using an <u>AvaSpec-2048 spectrometer</u> (Avantes, USA). 50 uM of CPR were loaded into 1mL cuvettes with a 1cm pathlength and measured prior to reduction with NADPH. Subsequent reductions were completed with increasing concentrations of NADPH adding, 12.5μM, 12.5μM, 25μM, 25μM and 25μM. The amount of NADPH added never exceeded 5% of the reaction volume. After each addition of NADPH, the reaction mixture was left to incubate for 1min before recording the spectra. Spectra were analyzed from 300-1,100 nm and recorded using the <u>AfterMath Electrochemical Studio</u> program version 1.6.10513 (Pine Research Instrumentation, USA).

### Catalytic Assays with Cytochrome c, NADPH, and NADH

Kinetic analyses of the two enzymes was performed using the <u>BioTek Synergy H1 plate reader</u> from Agilent Technologies (Santa Clara, CA) with black, non-binding, flat clear bottom 96-well plates from Greiner Bio-One (Monroe, NC). Reactions were run in 300μL volumes and contained: 10 nM of enzyme, buffer (50 mM Tris-HCl pH 7.5), and varying concentrations of substrates/ligands. For determining kinetics with cytochrome *c*, a concentration range of: 0.50 μM, 0.75 μM, 1 μM, 2 μM, 4 μM, 8 μM, 10 μM, 20 μM, 36 μM, and 50 μM was used. All reactions were initiated by adding 100 μM of NADPH. For determining kinetics with NADPH, a concentration range of: 1 μM, 2 μM, 4 μM, 10 μM, 20 μM, 50 μM, 75 μM, and 100 μM was used. All reactions were initiated by adding 10 μM of cytochrome *c*. For determining kinetics with NADH, a concentration range of: 200 μM, 500 μM, 1 mM, 2 mM, 5 mM, 10 mM, 20 mM, 30 mM, and 50 mM was used. All reactions were initiated by adding 10 μM of cytochrome *c*.

The reduction activity of enzymes was shown by the changes in absorption (ABS) of Cytochrome c recorded across 95s at 550 nm based on an extinction coefficient of 21/mM/cm for quantification (Guengerich et al., 2009). Initial-rate behaviour was monitored in real time for consistency and experiments were repeated as needed to resolve technical errors such as incorrect protein dilutions, pipetting errors, and bubbles in the reaction. All experiments included five technical replicates (five reaction wells) and three experimental replicates. Two biological replicates were included for AaCPR2.

### ***M***odel Assumptions/Statistical Analysis for Kinetics

Raw data from technical replicates and independent experimental repeats were pooled following quality control assessment (comparing initial velocities at the time of experimentation).

Replicates exhibiting clear technical artifacts, including abnormal initial absorbance values or initial slopes inconsistent with the distribution of replicate measurements, were excluded prior to analysis. Remaining replicates displaying consistent behavior across experimental repeats were treated as independent measurements for subsequent kinetic analysis. The Python code for organising the data and plotting Michaelis-Menten curves can be accessed through this <u>Github</u> <u>repository</u>. Initial velocities were determined from the maximal linear region (R² ≥ 0.98) for each trace and measured across replicates at each substrate concentration. Kinetic parameters were obtained by nonlinear regression of the Michaelis–Menten equation using weighted least squares.

The inverse of the experimental variance at each substrate concentration was used as the weighting factor. Parameter uncertainties were estimated from the covariance matrix of the fit and are reported as standard errors. Ninety-five percent confidence intervals were calculated assuming a Student’s *t* distribution. Plots were generated to show mean initial velocities ± standard deviation with the best-fit model for visualization.

### Differential Scanning Fluorimetry/Thermofluor Assay

Thermofluor assays were performed on a <u>Bio-Rad CFX96 real-time system C100 thermal cycler</u>, using 96-well RT-PCR plates (Bio-Rad, USA). Prior to running each plate in the thermal cycler, a 5 μL volume of 10x SYPRO dye was added to each reaction volume to ensure that fluorescent changes were recorded as the proteins unfolded. For general ligand testing, the concentrations of NADPH, cytochrome c, and NADH were increased 3-5x of their Km values to ensure optimal activity of the experimental system and used according to the following experimental design. For AaCPR1: No ligand, 30 μM NADPH, 20 μM cytochrome *c*, 7.5 mM NADH, 30 μM NADPH + 20μM cytochrome *c*, and 7.5 mM NADH + 20 μM cytochrome *c*. For AaCPR2: No ligand, 30 μM NADPH, 20 μM cytochrome *c*, 15 mM NADH, 30 μM NADPH + 20 μM cytochrome *c*, and 15 mM NADH + 20 μM cytochrome *c*.

To investigate the effect of increasing NADPH on enzyme thermostability with 20μM cytochrome *c*, an NADPH concentration range of 1μM, 2μM, 5μM, 10μM, 30μM, 100μM, and 200μM, was tested against AaCPR1 and AaCPR2. The same NADPH range was also tested without cytochrome *c* to investigate its effects on enzyme thermostability alone.

To investigate the effect of increasing cytochrome *c* on enzyme thermostability with 30 uM NADPH, a cytochrome *c* concentration range of 0.75μM, 1μM, 2μM, 5μM, 10μM, 20μM, 50μM, and 100μM was tested against AaCPR1 and AaCPR2. The same cytochrome *c* range was also tested without cytochrome *c* to investigate its effects on enzyme thermostability alone.

All experiments included no-ligand conditions as a baseline and enzyme free negative controls were carried out for each reaction condition, including, with dye only, NADPH + 20μM cytochrome *c* only, cytochrome *c* + 30μM NADPH only, NADPH only, and cytochrome *c* only.

All experiments described above were carried out as 50 μL reactions in each well. The reaction cycle held samples at 25°C for ten minutes and then increased in 0.5°C increments until 95°C. Each temperature increase was held for 5 s and the cycle was terminated after reaching 95°C. Tm values were calculated by the first derivative of the raw fluorescence data in the CFX software.

All experiments included three technical replicates and three experimental replicates. Two biological replicates were used for AaCPR2.

### Thermofluor Data Plotting and Analysis

The raw fluorescence data in relative fluorescence units (RFU) after each measurement was exported from the CFX software and used to run in-depth melt curve peak analyses in a customized Python tool <u>DSF_PeakSpeak</u>. Derivative melting curves were calculated as d(RFU)/dT using Savitzky–Golay smoothing. Prior to establishing parameters for melt peak calling, the negative controls were visualized alongside treatment derivative curves to exclude any fluorescence behaviour that was associated with ligand or dye related artefacts. A temperature range between 30°C and 75°C was established in the derivative analysis to exclude inaccurate peak calling resulting from protein aggregation at extreme low and high temperatures. Optimal peak calling was based on the plotted derivative curves and the standards established in the following description. Two transitions (T1, T2) were assigned based on temperature order, rather than energetic dominance, to provide a consistent labeling framework across all wells and conditions. Additionally, to reduce noise from minor fluctuations or aggregation artifacts, transitions with a low derivative magnitude (below a depth threshold of 21 a.u.) were excluded and only temperature values from the last two unfolding transitions per well were reported in the temperature tables.

Apart from derivative peak plotting, melting temperatures were also represented as normalized curves. These present a more simplified version of the melt peaks from the derivative data. For these plots, raw fluorescence data were normalised to a 0–1 scale before averaging across replicates. Melting temperatures were estimated from the maxima of the normalized curves.

These normalised melting temperatures are typically slightly higher than the derivative peak maxima (dRFU/dT) because derivative peaks correspond to the steepest slope rather than the absolute fluorescence plateau. Normalized curves are presented as supplementary validation of the trends observed in derivative curves.

Ordinary least squares (OLS) regression models were run across all Thermofluor datasets (including all replicates under each condition). The pooled data was used with OLS to determine the significance of NADPH and cytochrome *c* effects on the two enzymes. All ligand conditions came out as statistically significant.

### Protein Modeling

The <u>ColabFold container</u> was sourced from Github and run on the Jetstream2 high performance computer server from Indiana University. ColabFold was used to generate predictions based on multiple sequence alignment data for AaCPR1, AaCPR2, ATR1 and ATR2. The highest ranked generative models were used for AaCPR1 and AaCPR2. To develop reliable FAD, FMN and NAP binding sites, the proteins were aligned to the crystal structures of ATR2 (<u>5GXU</u>) and SbCPR2b (<u>7SV0</u>) for detection of conserved residues and protein geometry. Due to apparent distortion in the predicted FAD and NAP binding pockets of conserved tryptophan residues, the local structures of the AaCPRs were refined based on the homologous ATR2 and SbCPR2b crystal structures to restore canonical FAD and NAP stacking geometry via residue rotation based on the <u>Dunbrack Rotamer Library</u>. All visualisations, including initial ligand positioning predictions with <u>Foldseek</u> and hydrogen bond detection were done in <u>ChimeraX</u> from UCSF. Modelling accuracy was further improved by optimising residue contacts and reducing steric clashes with ligands through ligand repositioning.

## Supporting information

Replicate DSF Curves

## Acknowledgements

Thank you to the members of the Makris and Baryicki and Simpson labs in Molecular and Structural Biochemistry: Han Pham for his assistance with protein dialysis, Sydney Skirboll for her assistance with spectral experiments, Emily Allego and Brenna Zimmer for their assistance with Western blotting, enzyme kinetics, and Thermofluor.

Thank you to Dr. Imara Perrera for her interim advising and assistance in advancing the purification of AaCPR2.

This work used Jetstream2 at Indiana University through allocation [allocation number] from the Advanced Cyberinfrastructure Coordination Ecosystem: Services & Support (ACCESS) program, which is supported by National Science Foundation grants #2138259, #2138286, #2138307, #2137603, and #2138296 Molecular graphics and analyses performed with UCSF ChimeraX, developed by the Resource for Biocomputing, Visualization, and Informatics at the University of California, San Francisco, with support from National Institutes of Health R01-GM129325 and the Office of Cyber Infrastructure and Computational Biology, National Institute of Allergy and Infectious Diseases.

