## Supplementals S1-S7 for "Enzymatic and Biophysical Analysis of two Highly Related Cytochrome P450 Reductases from *Artemisia annua* Reveals Differences in Their Ligand Interactions and Domain Motions"

### Supplemental Data

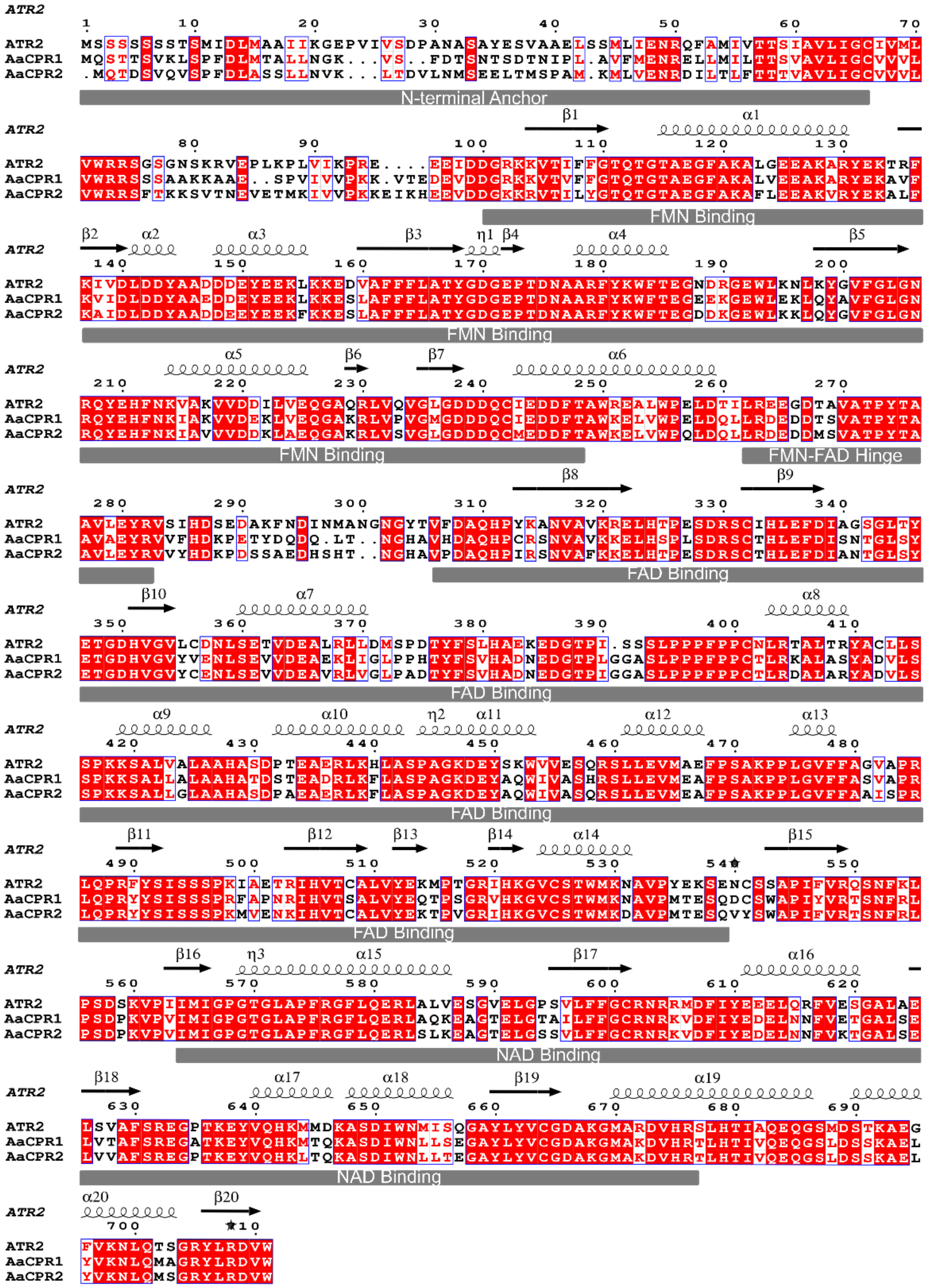

Supplemental Figure 1. Multi-sequence alignment of *Arabidopsis thaliana* CPR2 (ATR2), *Artemisia annua* CPR1 (AaCPR1) and *Artemisia annua* CPR2 (AaCPR2). Highlighted residues represent sequence identity (red) and sequence similarity (white). The respective binding domains are highlighted in grey: N-terminal anchor, FMN binding domain, FMN-FAD hinge, FAD binding domain, and NAD binding domain. Alignment is overlayed with the secondary structure of ATR2 (e.g., α helices and β sheets).

| Preparation/Vector | Primer-pairs | Thermal Conditions |
|---|---|---|
| AaCPR1 full length sequence | F: 5'-ATGCAATCAACAACTTC-3' | 98°C X 30" + 35(98°C X 10"+ 50°C X 30" + 72°C x 30") + 72°C X 5' |
| | R: , 5' TTACCATACATCACGGAGAT-3 | |
| AaCPR2 full length sequence | F: 5'-ATGCAAACAGATTCCG-3' | 98°C X 30" + 35(98°C X 10"+ 50°C X 30" + 72°C x 30") + 72°C X 5' |
| | R: 5' TTACCAAACATCGCGAAGATATCT-3' | |
| AaCPR1 truncated sequence | F: 5'- CACCAAGAAAGCGGCGGAGTCGCCG-3' | 98°C X 30" + 35(98°C X 10"+ 50°C X 30" + 72°C x 30") + 72°C X 5' |
| | R: 5' TTACCATACATCACGGAGATATCT-3' | |
| TEV site + truncated aaCPR2 in pENTR/D-TOPO | TEV-CPR2t-F-CACC: 5'CACCGAAAACCTGTATTTTCAGGGAAGGAGATCG TTTACTAAGAAATCGGT-3' | 98°C X 30" + 35(98°C X 10"+ 55°C X 30" + 72°C x 30") + 72°C X 5' |
| | CPR2-R: 5'-TTACCAAACATCGCG-3' | |

Table S1. Primer pairs and thermocycler conditions for PCR reactions to clone AaCPR1 pDEST17 and AaCPR2 HisMBP-pDEST17.

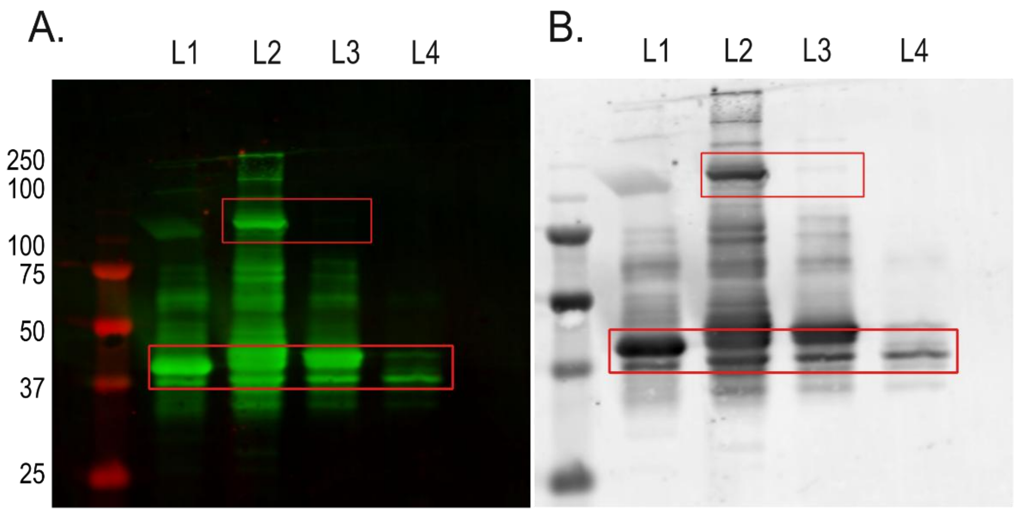


Supplemental Figure 2. Western blot with an MBP polyclonal antibody on His-MBP AaCPR2 purification fractions in (A) colour and (B) inverted to B&W. L1 = crude extract/cell lysate, L2 = elution 1 of the fusion protein from Ni-NTA column, L3 = TEV digestion of the fusion protein, L4 = Elution from the MBP affinity column. The fusion protein, MBP-AaCPR2 is ~139kDa and is visible in L1 and L2. After digestion, the fusion protein is cleaved leading to a loss of the 139kDa band in L3. The partial MBP protein itself is ~42kDa and is present in all lanes. L4 depicts the MBP protein after it has been separated from the cleaved AaCPR2.

| Buffer Type | pH |
|---|---|
| Citric acid | 5.0 |
| Citric acid | 5.5 |
| Potassium phosphate | 6.0 |
| Potassium phosphate | 6.5 |
| Potassium phosphate | 7.0 |
| Potassium phosphate | 7.5 |
| Potassium phosphate | 8.0 |
| Tris-HCl | 7.5 |
| Tris-HCl | 8.0 |
| Tris-HCl | 8.5 |
| Tris-HCl | 9.0 |

Table S2. A list of the tested buffers and their respective pH for optimal kinetic performance of AaCPR1 and AaCPR2.

A. NADPH Dependency + 20uM Cytochrome c

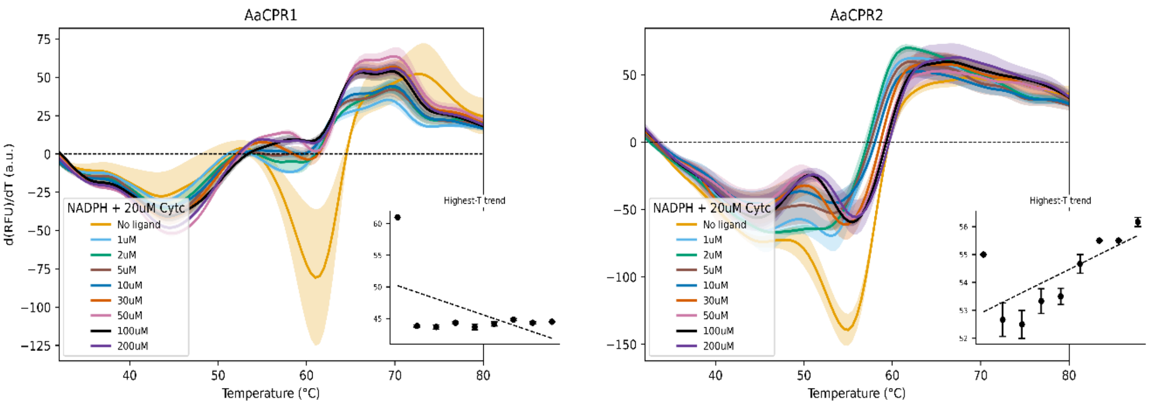


| Condition | Melting Temp(s) °C |
|---|---|
| No ligand | 43.8 ± 0.2 \| 61.0 ± 0.0 |
| 1uM | — \| 43.8 ± 0.2 |
| 2uM | 37.0 \| 43.7 ± 0.3 |
| 5uM | 36.0 \| 44.3 ± 0.2 |
| 10uM | — \| 43.7 ± 0.4 |
| 30uM | 37.5 \| 44.2 ± 0.3 |
| 50uM | 37.0 \| 44.8 ± 0.2 |
| 100uM | 36.5 \| 44.3 ± 0.2 |
| 200uM | — \| 44.5 ± 0.0 |

| Condition | Melting Temp(s) °C |
|---|---|
| No ligand | 44.5 \| 55.0 ± 0.0 |
| 1uM | 46.2 ± 0.9 \| 52.7 ± 0.6 |
| 2uM | 47.2 ± 1.0 \| 52.5 ± 0.5 |
| 5uM | 45.0 ± 1.3 \| 53.3 ± 0.4 |
| 10uM | 44.5 ± 0.5 \| 53.5 ± 0.3 |
| 30uM | 44.7 ± 0.3 \| 54.7 ± 0.3 |
| 50uM | 44.2 ± 0.2 \| 55.5 ± 0.0 |
| 100uM | 44.8 ± 0.2 \| 55.5 ± 0.0 |
| 200uM | 44.5 ± 0.0 \| 56.2 ± 0.2 |

B. Cytochrome c Dependency + 30uM NADPH

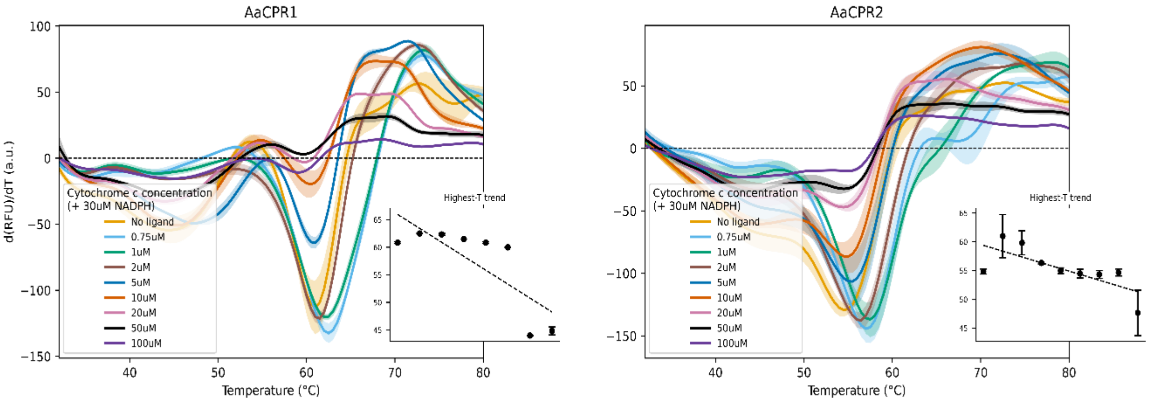


| Condition | Melting Temp(s) °C |
|---|---|
| No ligand | 44.2 ± 0.6 \| 60.8 ± 0.2 |
| .75uM Cytochrome c | — \| 62.5 ± 0.0 |
| 1uM Cytochrome c | — \| 62.3 ± 0.2 |
| 2uM Cytochrome c | — \| 61.5 ± 0.0 |
| 5uM Cytochrome c | 47.8 ± 0.6 \| 60.8 ± 0.2 |
| 10uM Cytochrome c | 45.3 ± 0.7 \| 60.0 |
| Cytochrome c | 36.5 \| 44.0 ± 0.0 |
| 50uM Cytochrome c | 43.0 \| 44.8 ± 0.7 |

| Condition | Melting Temp(s) °C |
|---|---|
| No ligand | — \| 54.8 ± 0.4 |
| .75uM Cytochrome c | 56.5 \| 61.0 ± 3.8 |
| 1uM Cytochrome c | 51.0 ± 6.0 \| 59.8 ± 2.1 |
| 2uM Cytochrome c | 44.5 \| 56.3 ± 0.2 |
| 5uM Cytochrome c | 49.0 \| 55.0 ± 0.5 |
| 10uM Cytochrome c | 45.5 ± 1.0 \| 54.5 ± 0.8 |
| Cytochrome c | 45.3 ± 0.2 \| 54.3 ± 0.7 |
| 50uM Cytochrome c | 44.0 ± 0.3 \| 54.7 ± 0.6 |
| 100uM Cytochrome c | 43.5 \| 47.7 ± 3.9 |

Figure S3. Melting temperature peaks for AaCPR1 and AaCPR1. (A) Increasing NADPH concentrations (1-200μM) with 20μM cytochrome *c.* AaCPR1 loses its second melting transition when both NADPH and 20μM cytochrome *c* are applied. The first melting transition does not change when NADPH is added and increased. AaCPR2 maintains its first and second melting transitions, although the segregation between these two peaks does not become apparent until more than 10μM of NADPH is added. The temperature of the second melting peak drops below the no ligand $Tm_2$ baseline temperature when 1μM of NADPH is added, but then gradually increases as NADPH concentration increases. (B) Increasing cytochrome *c* concentrations (0.75-50/100μM) with 30μM NADPH. The addition of 0.75-2μM of cytochrome *c* to AaCPR1 results in a slight increase in melting temperature when compared to the no-ligand control. When cytochrome *c* is greater than 5μM, the second melting temperature for AaCPR1 decreases and the second transition is lost. The presence of the first melting transition in AaCPR2 is not very deep when compared to AaCPR1. When 0.75-5μM of cytochrome *c* is added, the second melting transition in AaCPR2 slightly increases. At 10μM of cytochrome *c* and above, the second melting transition drops to the same temperature as the ligand control. For both AaCPR1 and AaCPR2, energetic dominance is lost in the first and second melting transitions as cytochrome *c* increases.

A. Ligand Test Normalized Curves

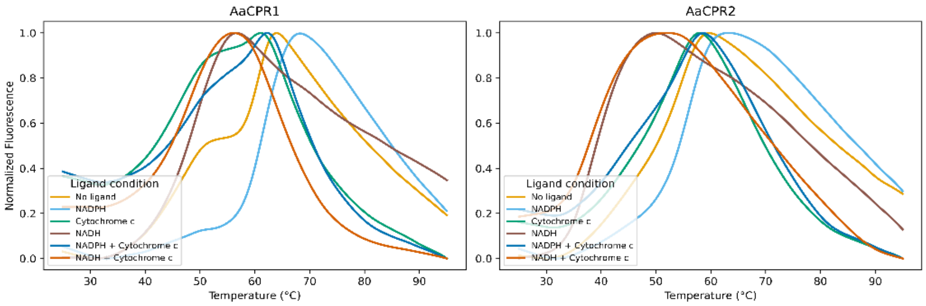


| Condition | Melting Temp(s) °C |
|---|---|
| No ligand | 64.0 |
| NADPH | 68.0 |
| Cytochrome c | 61.0 |
| NADH | 56.5 |
| NADPH + Cytochrome c | 62.0 |
| NADH + Cytochrome c | 56.0 |

| Condition | Melting Temp(s) °C |
|---|---|
| No ligand | 59.5 |
| NADPH | 63.5 |
| Cytochrome c | 58.0 |
| NADH | 50.0 |
| NADPH + Cytochrome c | 58.5 |
| NADH + Cytochrome c | 52.0 |

B. NADPH Dependency Normalized Curves

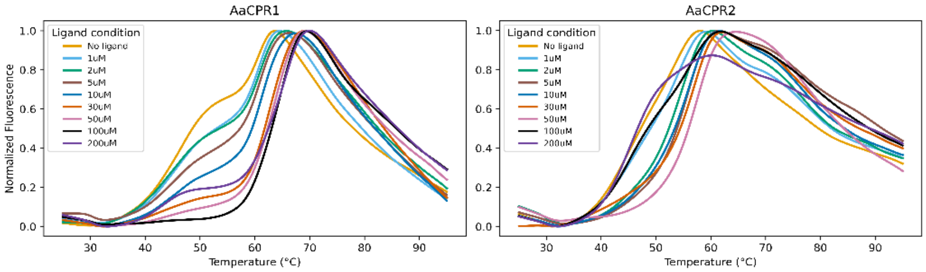


| Condition | Melting Temp(s) °C |
|---|---|
| No ligand | 63.5 |
| 1uM | 64.5 |
| 2uM | 66.0 |
| 5uM | 66.5 |
| 10uM | 67.5 |
| 30uM | 69.0 |
| 50uM | 69.0 |
| 100uM | 69.5 |
| 200uM | 70.0 |

| Condition | Melting Temp(s) °C |
|---|---|
| No ligand | 58.0 |
| 1uM | 59.0 |
| 2uM | 60.5 |
| 5uM | 61.5 |
| 10uM | 61.5 |
| 30uM | 62.0 |
| 50uM | 64.5 |
| 100uM | 62.0 |
| 200uM | 60.0 |

C. Cytochrome c Dependency Normalized Curves

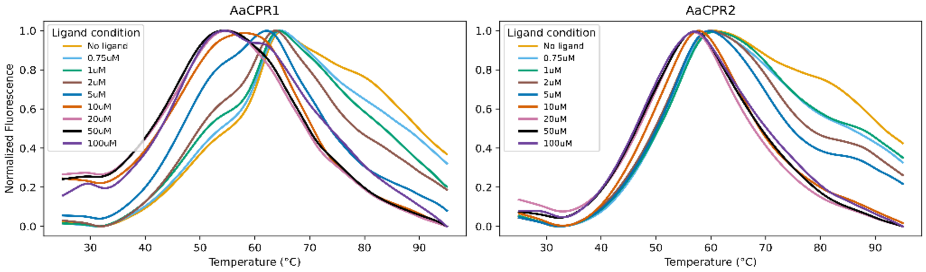


| Condition | Melting Temp(s) °C |
|---|---|
| No ligand | 65.0 |
| 0.75uM | 65.0 |
| 1uM | 64.5 |
| 2uM | 63.5 |
| 5uM | 62.0 |
| 10uM | 58.0 |
| 20uM | 55.0 |
| 50uM | 54.5 |
| 100uM | 54.5 |

| Condition | Melting Temp(s) °C |
|---|---|
| No ligand | 60.5 |
| 0.75uM | 60.0 |
| 1uM | 60.5 |
| 2uM | 60.0 |
| 5uM | 59.5 |
| 10uM | 58.0 |
| 20uM | 57.0 |
| 50uM | 57.0 |
| 100uM | 57.0 |

igure S4. Normalised curves for the ligand test panel (A), NADPH dependency (B), and cytochrome *c* dependency experiments (C). Normalised curves depict only the major second melting transition across all conditions; the temperature trends from the derivative curves however remain the same for this melting peak.

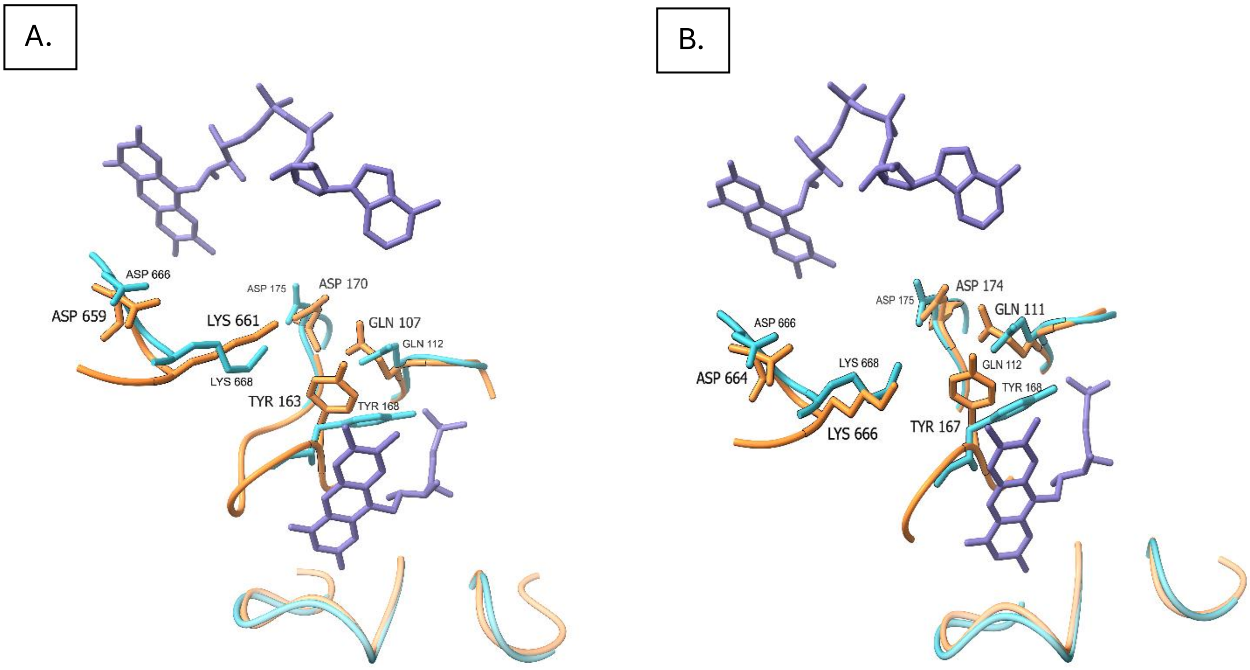


Fig. S5. Interfacing residues between the FAD and FMN binding domains. FAD (top) and FMN (bottom) co-enzymes are coloured in purple (A) Conserved residues between ATR2 (cyan) and AaCPR1 (orange) between key interfacing loops at the FAD and FMN binding domains. (B) Conserved residues between ATR2 (cyan) and AaCPR2 (orange) between key interfacing loops at the FAD and FMN binding domains.

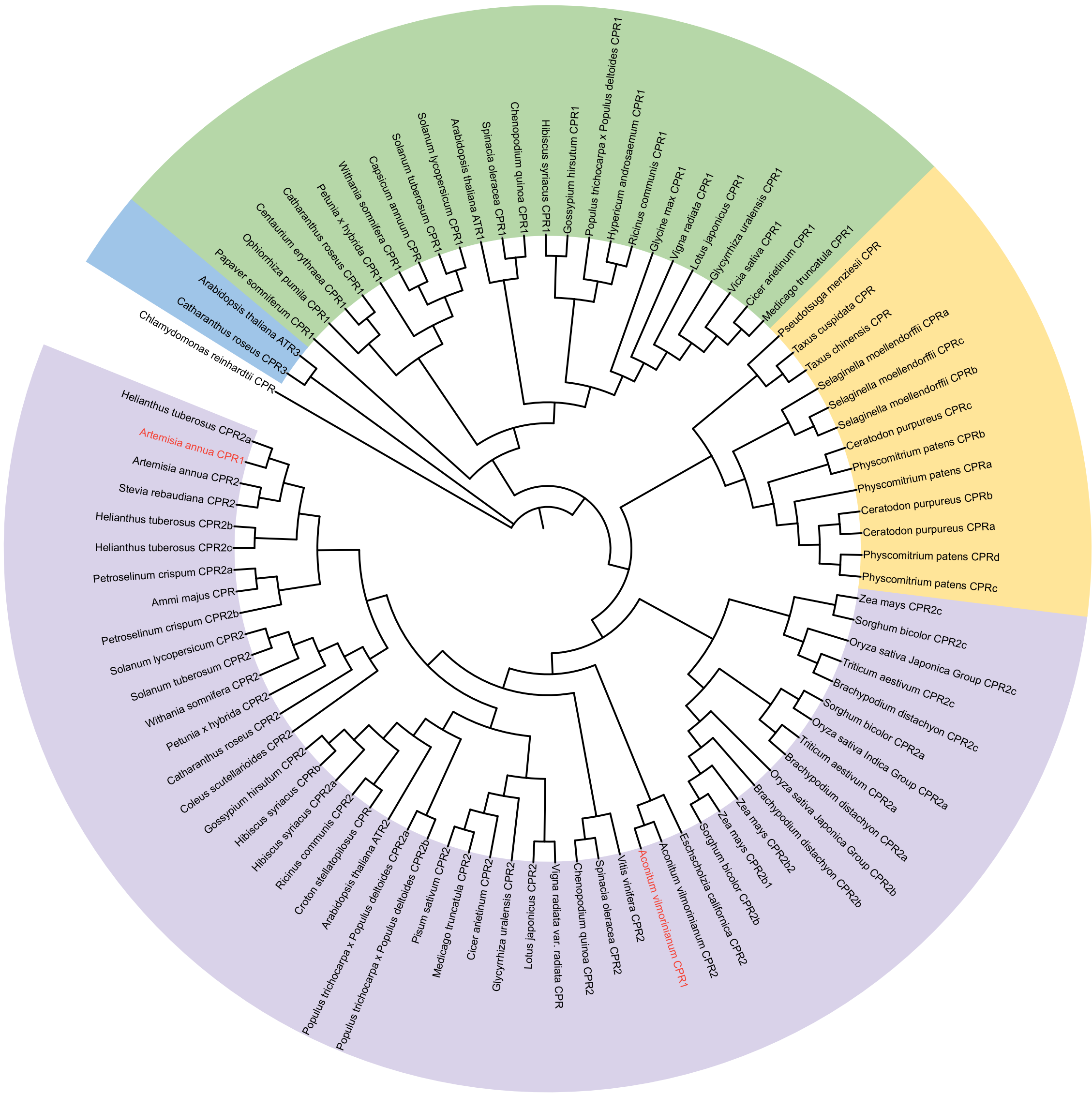


Figure S6. Phylogenetic tree of 89 CPRs including AaCPR1 and AaCPR2. Tree was built using the maximum likelihood method and bootstrapped with 100 iterations (Mega 11). *Chlamydomonas reinhardtii* was used as an outgroup (uncoloured). Notable clades are coloured: unique CPR3 enzymes which are Novel Reductase 1 (NR1) related (blue), CPR1 enzymes (green), Non-angiosperm CPRs (yellow), CPR2 enzymes (purple). *Artemisia annua* CPR1 and *Aconitum vilmorinianum* CPR1 text are highlighted in red due to their placement in the CPR2 clade.

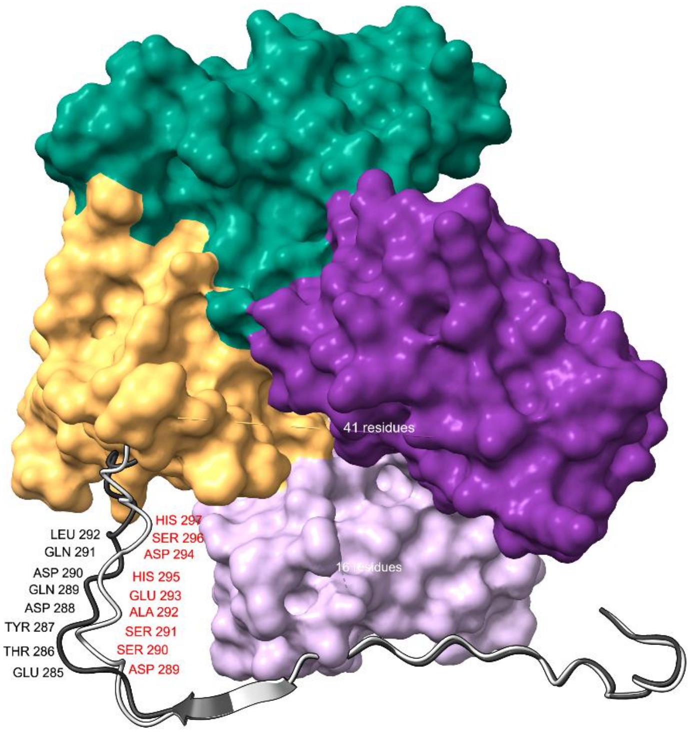


Figure S7. 3-dimensional surface of ATR2 from *Arabidopsis thaliana* with hinge-connecting domain segments from AaCPR1 and AaCPR2 superimposed (in grey). The highly variable residues of the connecting domain are highlighted between AaCPR1 (black text) and AaCPR2 (red text).
