## Supplementary material for "Enzymatic and Biophysical Analysis of two Highly Related Cytochrome P450 Reductases from *Artemisia annua* Reveals Differences in Their Ligand Interactions and Domain Motions": Replicate DSF Curves

A. Ligand Test CPR1

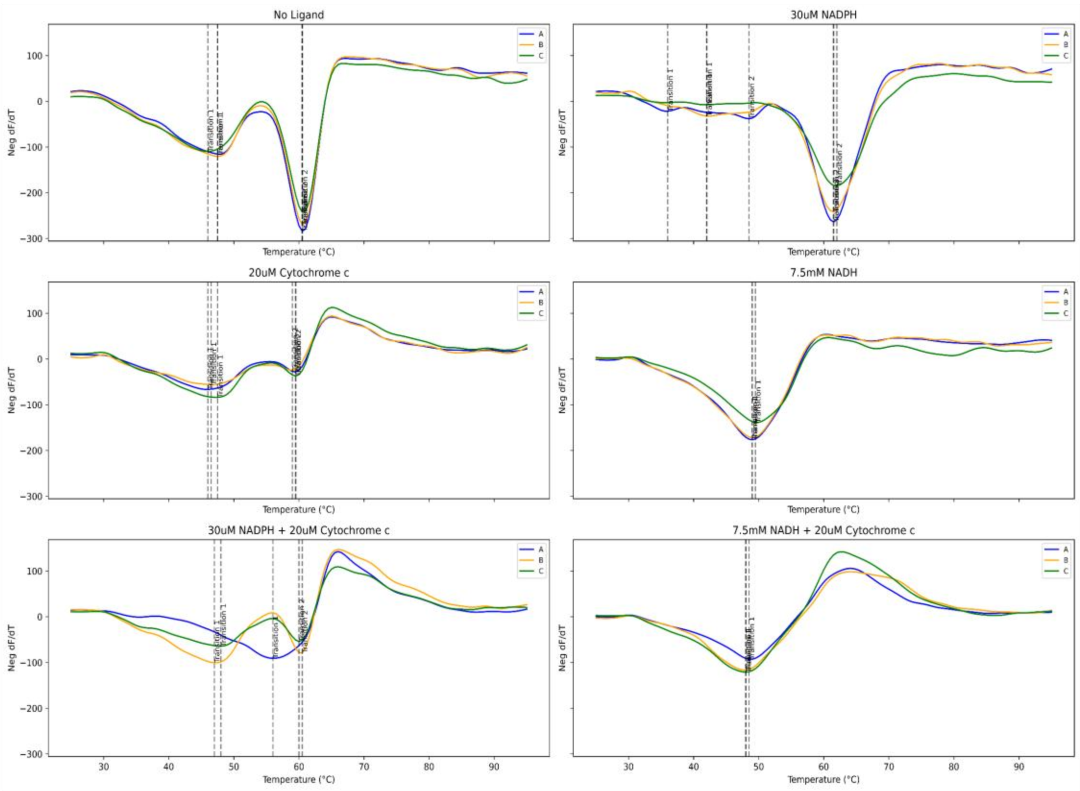


B. Ligand Test CPR2

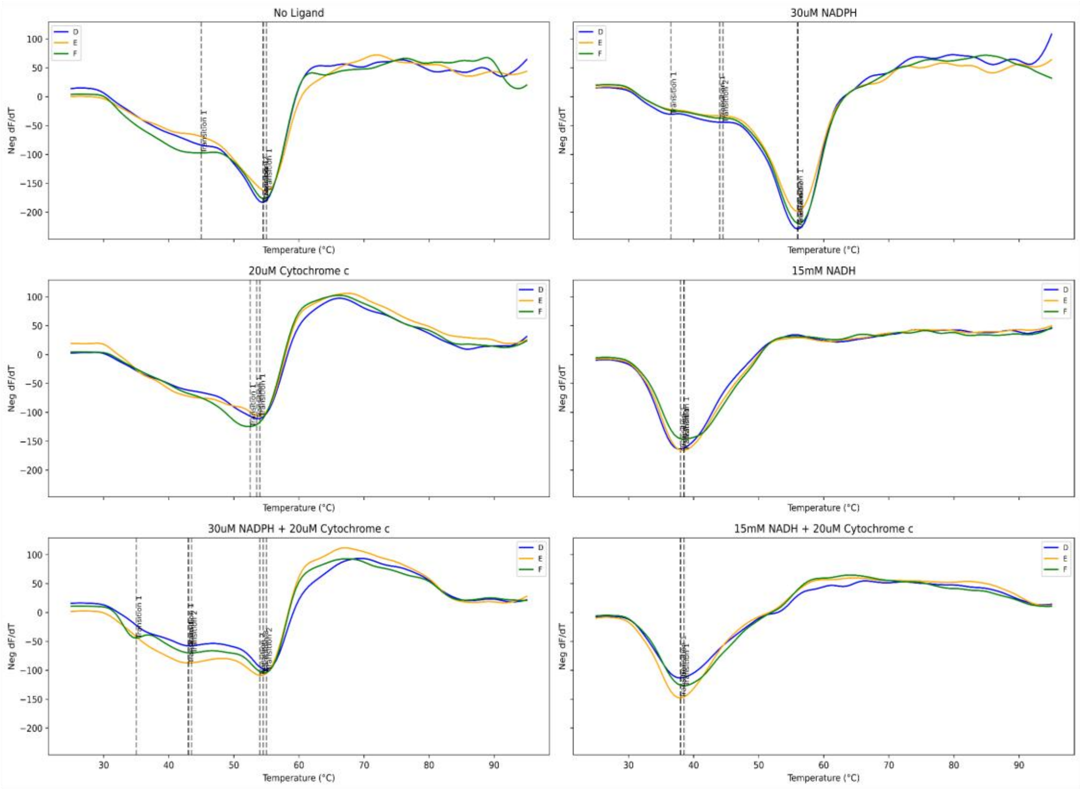

A. NADPH Test CPR1

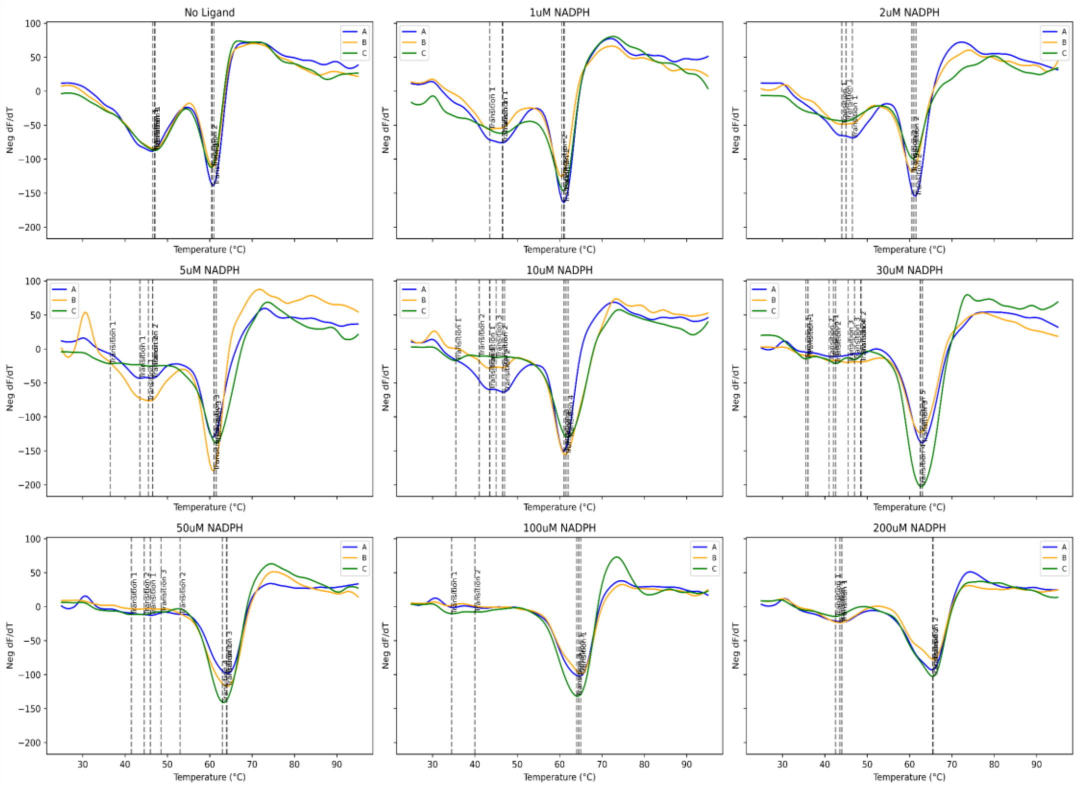


B. NADPH Test CPR2

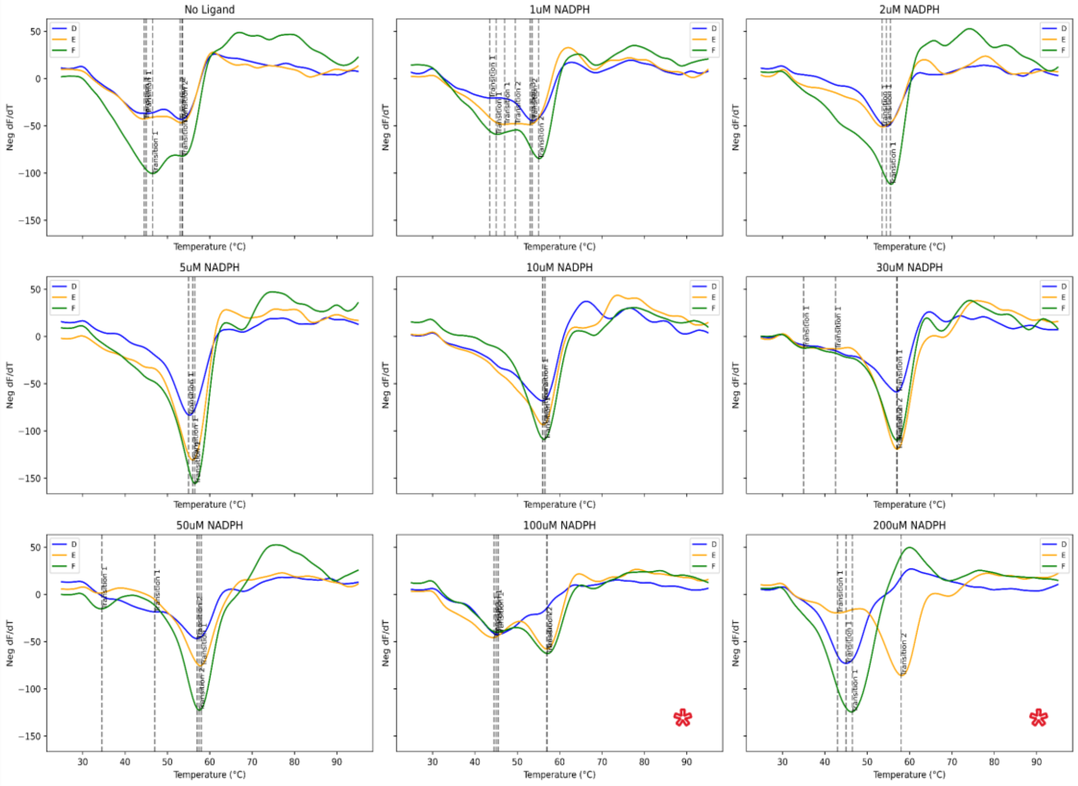

A. Cytochrome c Test CPR1

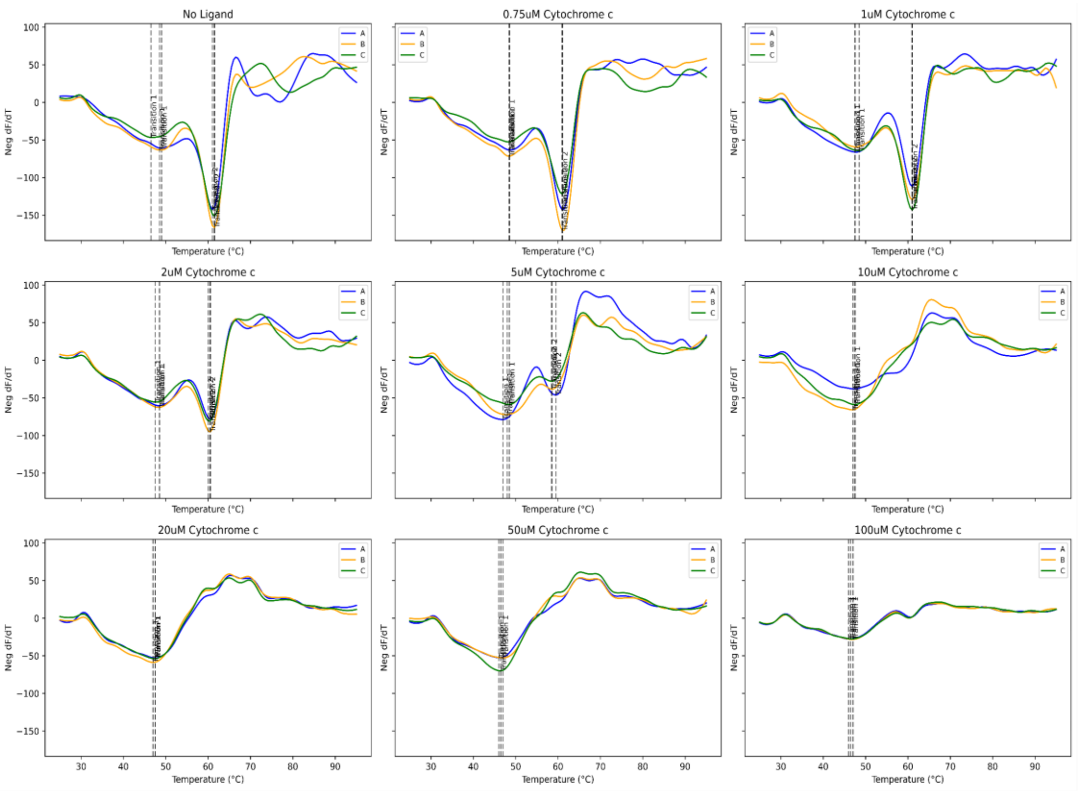


B. Cytochrome c Test CPR2

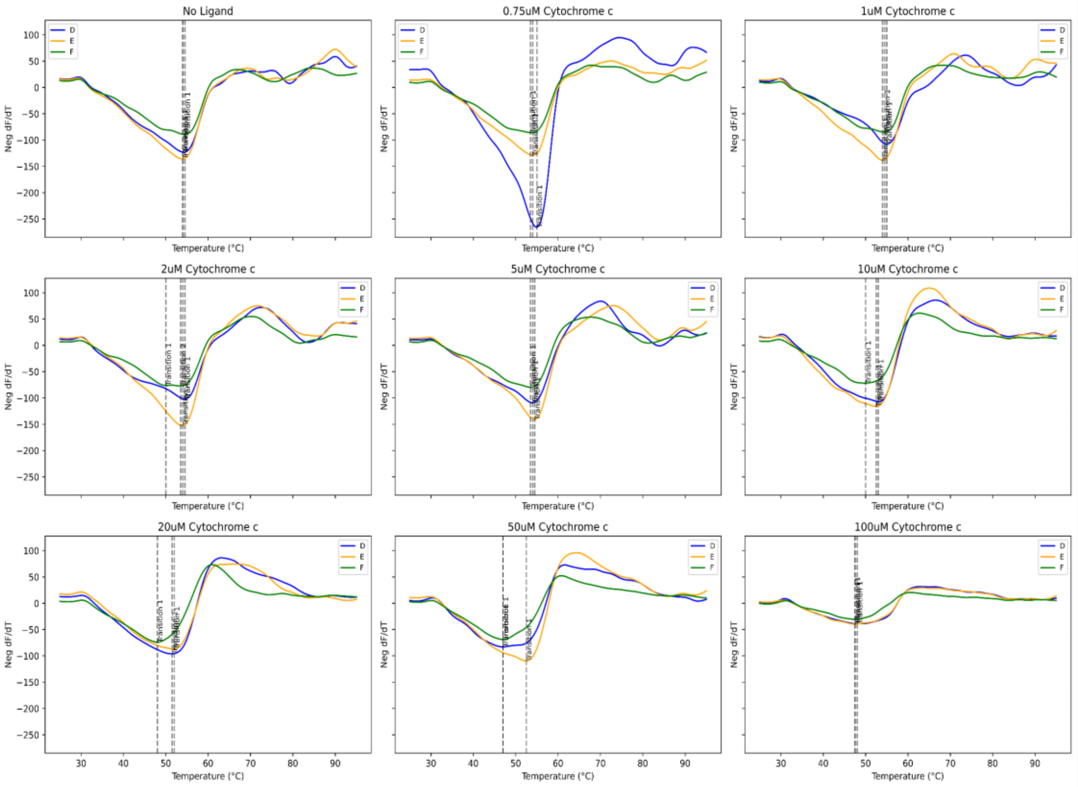

A. NADPH + 20uM Cyt c CPR1

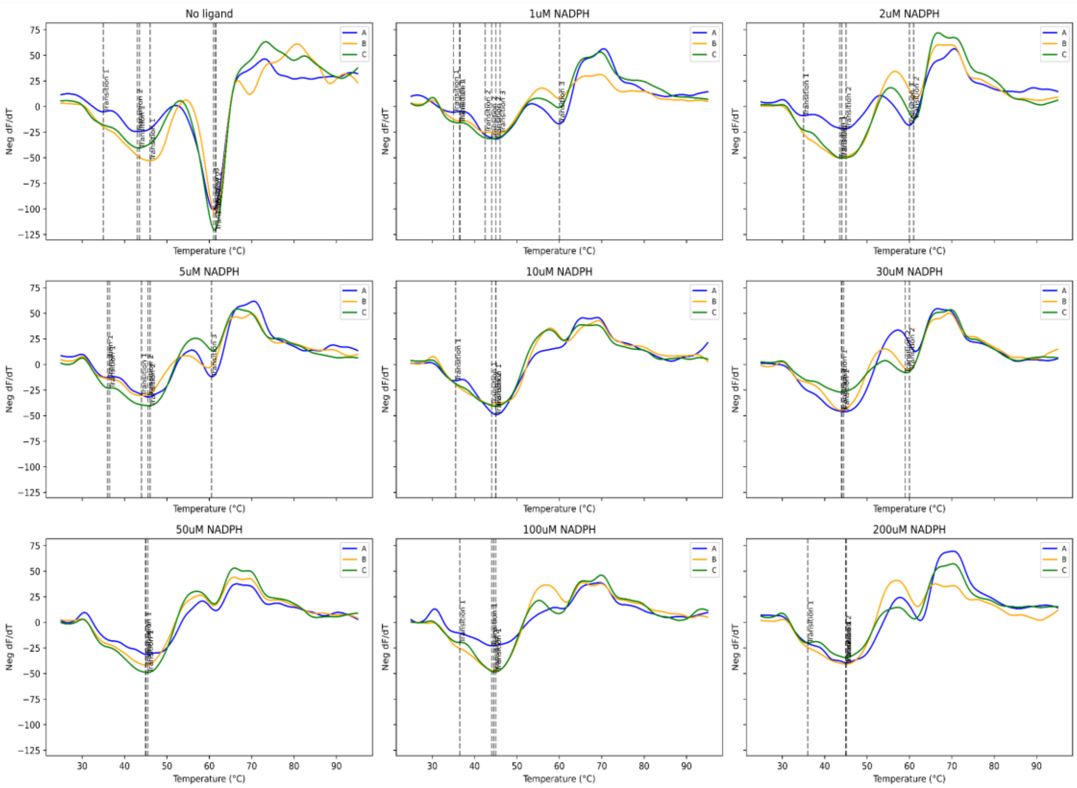


B. NADPH + 20uM Cyt c CPR2

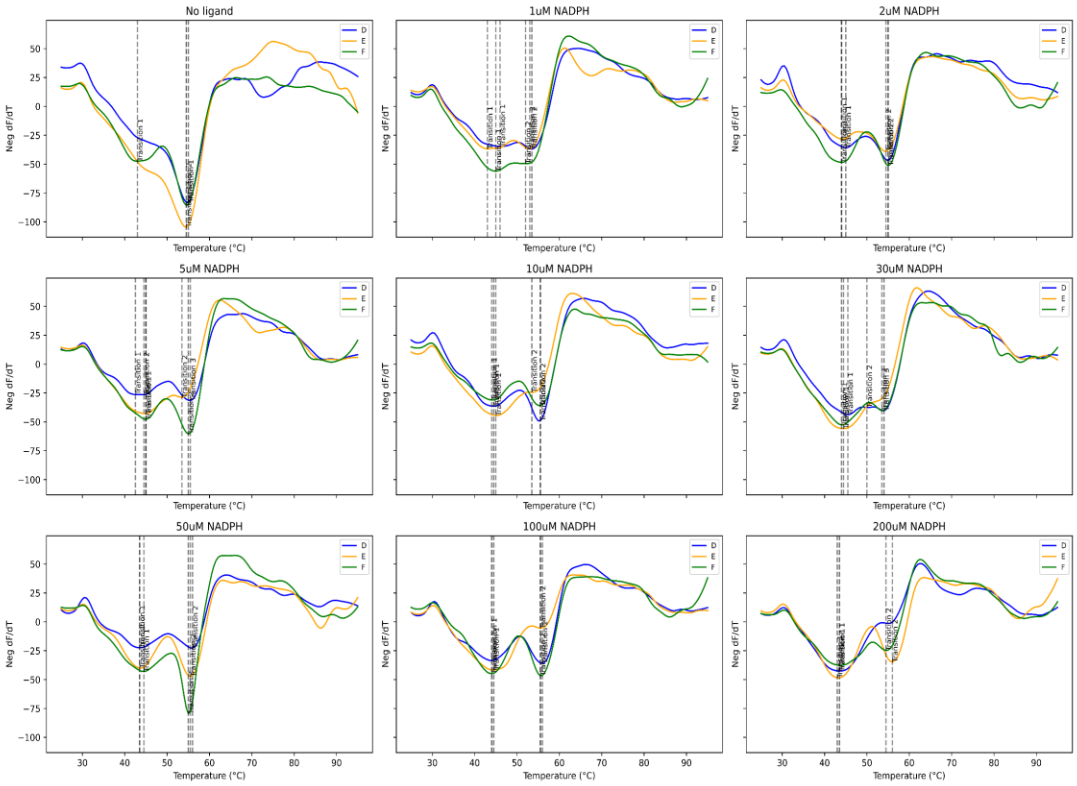

### A. Cyt c + 30uM NADPH CPR1

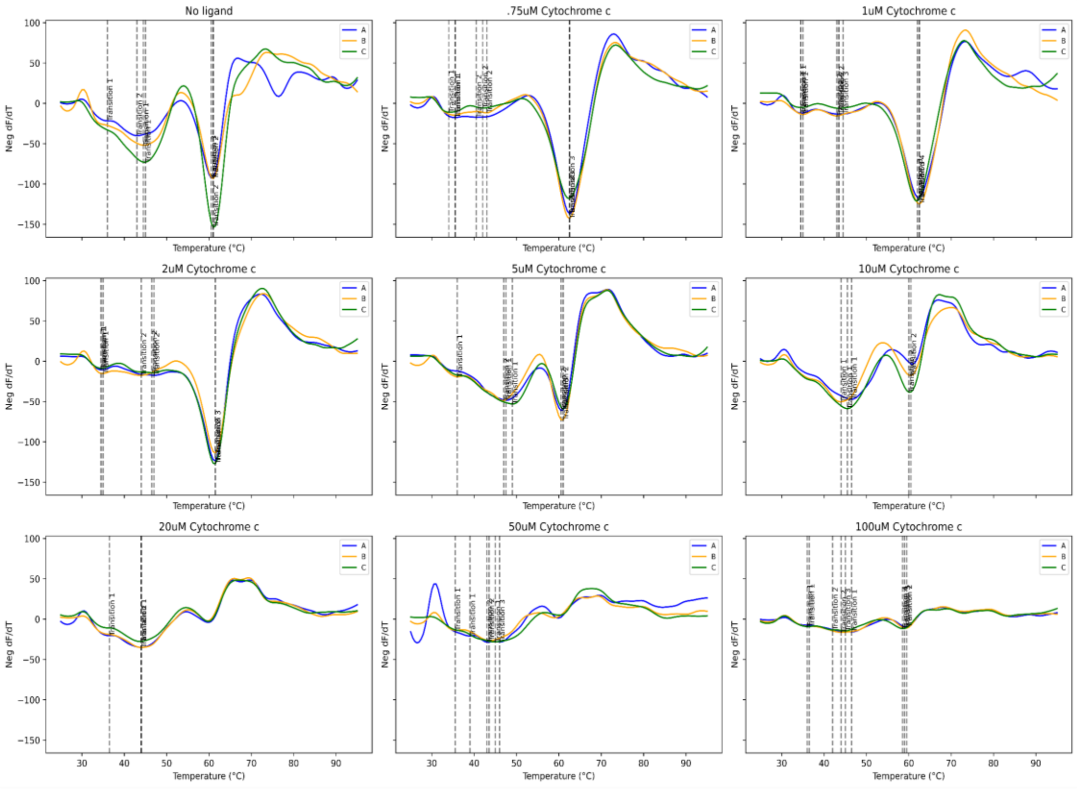


### B. Cyt c + 30uM NADPH CPR2

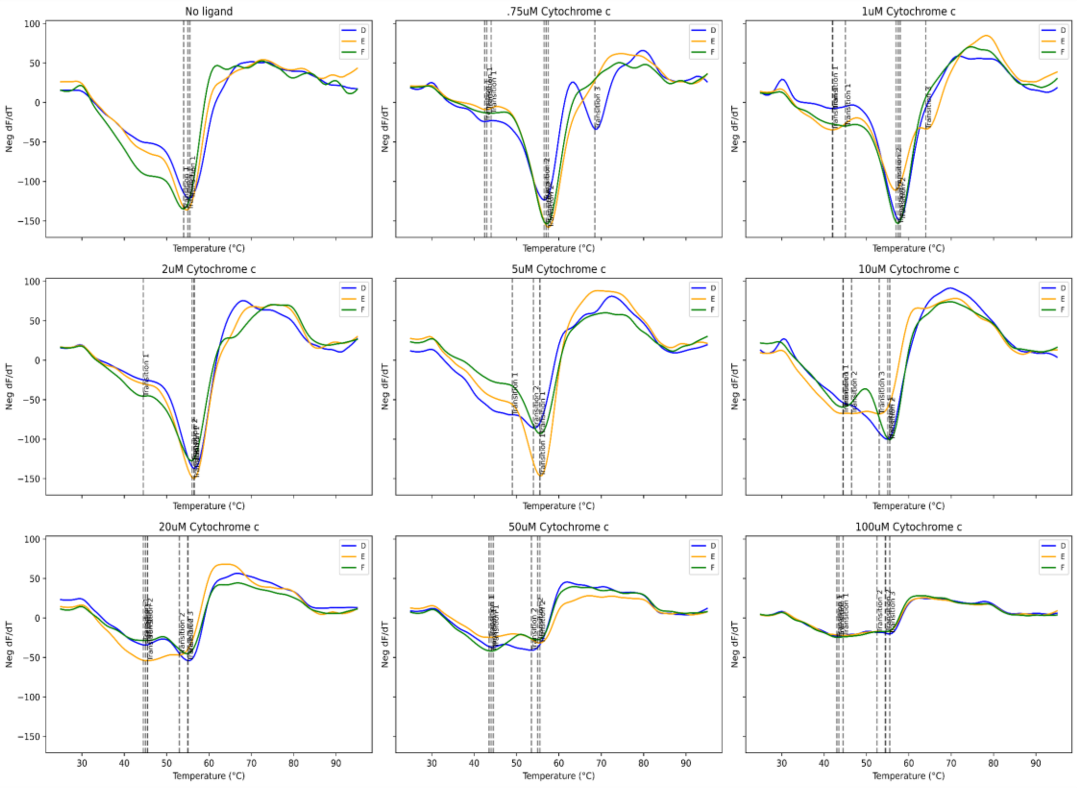
